# Proteobacteria drive significant functional variability in the human gut microbiome

**DOI:** 10.1101/056614

**Authors:** Patrick H. Bradley, Katherine S. Pollard

**Affiliations:** Gladstone Institutes, San Francisco, CA.; Division of Biostatistics, Institute for Human Genetics, and Institute for Computational Health Sciences, University of California, San Francisco, CA.

**Keywords:** human gut microbiome, Proteobacteria, Bacteroidetes, Firmicutes, variance, shotgun metage-nomics, statistical methods, functional redundancy, enterotypes

## Abstract

While human gut microbiomes vary significantly in taxonomic composition, biological pathway abundance is surprisingly invariable across hosts. We hypothesized that healthy microbiomes appear functionally redundant due to factors that obscure differences in gene abundance across hosts. To account for these biases, we developed a powerful test of gene variability, applicable to shotgun metagenomes from any environment. Our analysis of healthy stool metagenomes reveals thousands of genes whose abundance differs significantly between people consistently across studies, including glycolytic enzymes, lipopolysaccharide biosynthetic genes, and secretion systems. Even housekeeping pathways contain a mix of variable and invariable genes, though most deeply conserved genes are significantly invariable. Variable genes tend to be associated with Proteobacteria, as opposed to taxa used to define enterotypes or the dominant phyla Bacteroidetes and Firmicutes. These results establish limits on functional redundancy and predict specific genes and taxa that may drive physiological differences between gut microbiomes.

**Impact Statement:** A statistical test for gene variability reveals extensive functional differences between healthy humanmicrobiomes.

## 1 Background

The microbes that inhabit the human gut encode a wealth of proteins that contribute to a broad range of biological functions, from modulating the human immune system [1, 2, 3] to participating in metabolism [4, 5]. Shotgun metagenomics is revolutionizing our ability to identify protein-coding genes from these microbes and associate gene levels with disease [6], drug efficacy [7] or side-effects [8], and other host traits. For instance, gut microbiota associated with a traditional high-fiber agrarian diet encoded gene families involved in cellulose and xylan hydrolysis, which were absent in age-matched controls eating a typical Western diet [9]. The functional capabilities of the gut microbiome go beyond statistical associations; a number of microbial genes have now been causally linked to host physiology. Examples include the colitis-inducing cytolethal distending toxins of *Helicobacter hepaticus* [10] and the enzymes of commensal bacteria that protect against these toxins by producing anti-inflammatory polysaccharide A [11].

It is therefore surprising that healthy human gut microbiomes have been characterized as functionally stable (i.e., invariable), with largely redundant gene repertoires in different hosts. Several lines of evidence support this conclusion. First, biological pathway abundance tends to be less variable across metagenomes than it is between isolate genomes [12], suggesting strong selection for microbes that encode functions necessary for adaptation to the gut environment. Second, the relative abundances of pathways are strikingly invariable compared to the relative abundances of bacterial phyla in the same metagenomes [13, 12]. Thus, it appears that humans harbor phylogenetically distinct gut communities that all do more or less the same things, except in the context of disease or other extreme host phenotypes.

Functional redundancy deserves a closer look, however, because physiologically meaningful differences in gene abundances between healthy human microbiomes could easily have been missed. One primary factor may be that prior work did not look at quantitative abundances of individual genes, but instead mainly summarized function at the level of Clusters of Orthologous Groups (COG) categories and KEGG modules [13, 12, 14]. Summarizing genes into groups will not have power to detect one component of a pathway or protein complex that varies in abundance across hosts if other components are less variable. This masking of variable genes is likely to occur because the presence and abundance of most COG categories and KEGG modules will be dominated by core components (i.e., housekeeping genes) that are widely distributed across the tree of life and abundant in metagenomes. The only previous analyses of individual genes asked whether they were universally detected across all individuals sampled [12, 14]; however, universally-detected genes may still vary substantially in abundance, and conversely, lower-abundance invariable genes may not be universally detected merely due to sampling. This approach is also sensitive to read depth [12] and sample size [14]. Based on these observations, we were motivated to quantitatively investigate functional redundancy at the level of individual gene families.

To enable high-resolution, quantitative analysis of functional stability in the microbiome, we developed a statistical test that identifies individual gene families whose abundances are either significantly variable or invariable across samples. Our method incorporates solutions to three major challenges to studying functional redundancy with shotgun metagenomics data. The first key innovation of our approach is using a test statistic that captures residual variability after accounting for overall gene abundance. This modeling choice is important because abundant genes will be variable just by chance due to the correlation between mean and variance in any sequencing experiment. Conversely, phylogenetically restricted genes will have relatively low variance due to being less abundant. Furthermore, gene abundances can be sparse (i.e., zero in many samples). For all of these reasons simply ranking genes based on their variances would yield many false positives and false negatives.

A second benefit of our modeling approach is that we can adjust for systematic differences in a gene’s measured level between studies to allow for quantitative integration of data from multiple sources. Meta-analysis is essential for gaining sufficient power to detect variable genes across the range of mean abundance levels. It also ensures robustness and generalizability of discovered inter-individual differences, which occur by chance in small sets of metagenomes.

Finally, our method does not require predefined cases and controls, but instead enables discovery of genes that drive functional differences between microbiomes without prior knowledge of which groups of samples to compare. This is critical for the current phase of microbiome research, when many drivers of microbial community composition are unknown. Gene families that contribute to survival in one particular type of healthy gut environment should emerge as variable between hosts and their functions may point to drivers of community composition, mechanisms of microbe-host interactions, and biomarkers of presymptomic disease (e.g., pre-diabetes).

We applied our test to healthy gut metagenomes (n = 123) spanning three different shotgun sequencing studies and found both significantly invariable (3,768) and variable (1,219) gene families (FDR<5%). Many pathways, including some commonly viewed as housekeeping or previously identified as invariable across gut microbiota (e.g., central carbon metabolism and secretion), included significantly variable gene families. Phylogenetic distribution (PD) correlated overall with variability in gene family abundance, and exceptions to this trend highlight functions that may be involved in adaptation, such as two-component signaling and specialized secretion systems. Finally, we show that Proteobacteria, and not the major phyla Bacteroidetes and Firmicutes, are a major source for genes with the greatest variability in abundance across hosts, suggesting a relationship between inflammation and gene-level differences in gut microbial functions. This approach to discovering functions that distinguish microbial communities is applicable to any body site or environment.

## 2 Results

### 2.1 A new test captures the variability of microbial gene families

We present a model that enables gene family abundance to be quantitatively compared across metagenomes for thousands of microbial genes. In shotgun metagenomics data, different gene families vary widely in average abundance (Figure 1). Gene family abundances can also vary by study, both because of biological differences between populations, and for technical reasons including library preparation, amplification protocol, and sequencing technology (see, e.g., Figure 1 G-H). To account for such effects, we fit a linear model of log abundance *D*_*g,s*_ for gene *g* in sample s as a function of the overall mean abundance *µ*_*g*_ and a term *β*_*g,y*_ that quantifies the offset for each study *y:*

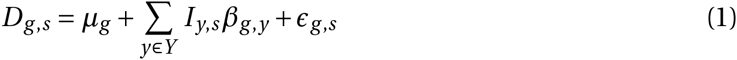

where *I*_*y,s*_ is an indicator variable that is 1 if sample *s* belongs to study y and 0 otherwise.

**Figure 1:**
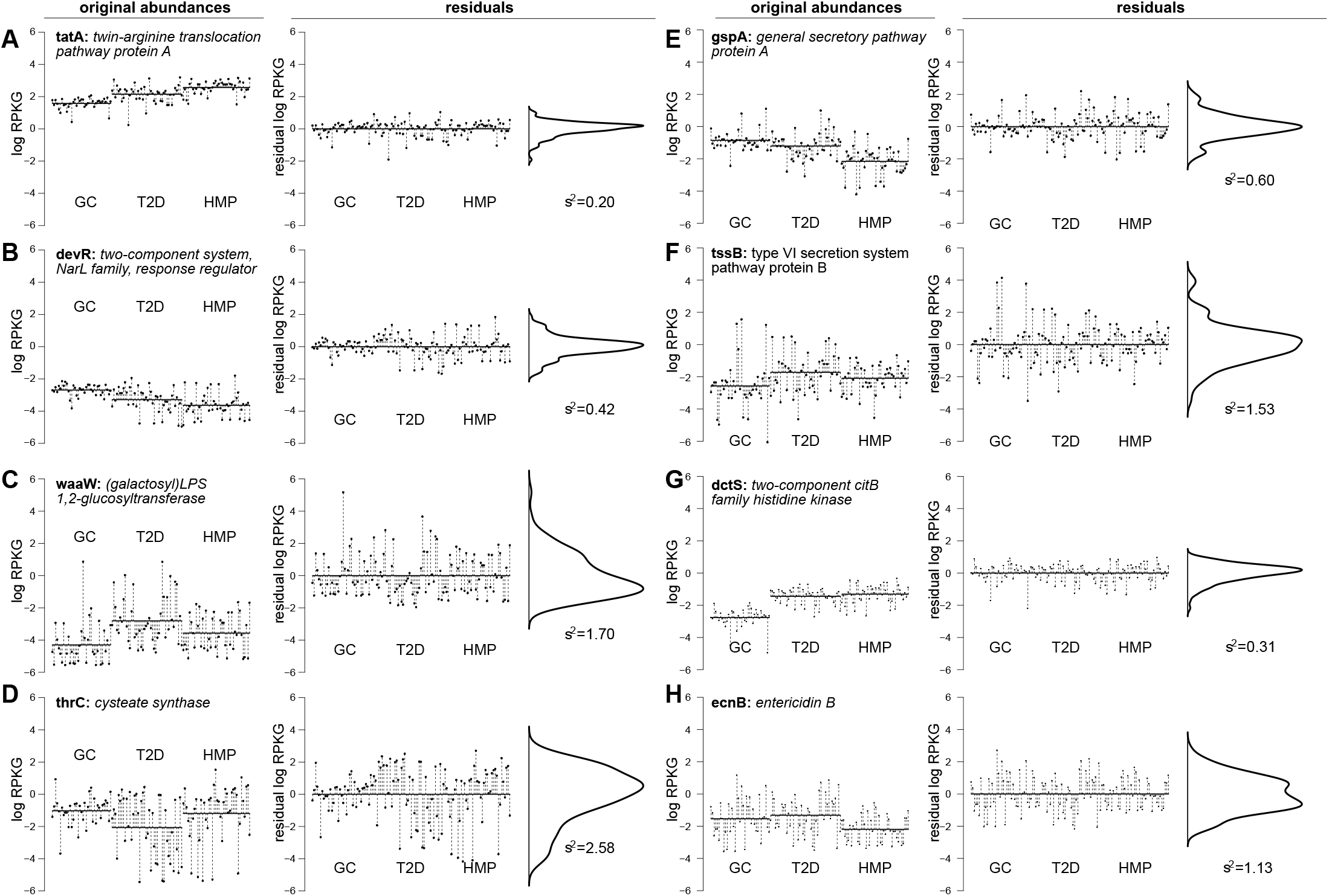
The residual variance statistic captures variation in gene families after accounting for between-study variation. Left panels (“original abundances”) show filled circles representing log-RPKG abundances for gene families from the KEGG Orthology (KO), with per-study means shown in solid horizontal lines and the distance from these means shown as dashed vertical lines. Right-hand panels (“residuals”) show the same gene families after fitting a linear model that accounts for these per-study means, with an accompanying density plot showing the distribution of these residuals. 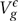 values in bold underneath density plots are the calculated variances of these residuals. These gene families are sets of orthologs corresponding to the genes A) tatA, B) devR, c) waaW, d) thrC, E) gspA, F) tssB, G) dctS, and H) ecnB. Panels A-B show two invariable gene families with relatively high (A) and low (B) average abundance; similarly, panels C-D show two variable gene families with relatively low (C) and high (D) relative abundances. Panels E-F show two gene families involved in secretion with similar abundances, but low (E) vs. high (F) variability. Finally, panels G-H show that both invariable (G) and variable (H) gene families can have substantial study-specific effects. (All gene families displayed were significantly (in)variable using the test we present, FDR ≤ 5%.)

The residual *ε*_*g,s*_ quantifies how much the abundance of gene *g* in sample *s* differs from the average abundance across samples in the same study as *s.* We denote the variance of the residuals across samples by 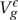. When this statistic is small, the gene has similar abundance across samples after accounting for study effects. A large value of 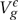 indicates that samples have very different abundances.

To assess the statistical significance of gene family variability, we compare the residual variance 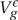 to a data-driven null distribution based on the negative binomial distribution (Figure 1—figure supplement 1, Methods). This approach is necessary because there is no straightforward formula for the p-value of 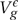. Our method looks for deviations from the null hypothesis that gene families in the dataset have the same mean-variance relationship (i.e., the same overdispersion). This choice of null is very important: if we were instead to simply test for high variance, regardless of mean abundance, highly abundant gene families (e.g., single-copy proteins in the bacterial ribosome) would be significantly variable despite being nearly universally present at equal abundance in each bacterial genome, because genes with high mean abundance would have high variance in any sequencing experiment. Meanwhile, thousands of lower-abundance gene families would appear to be significantly invariable simply by virtue of having relatively low read counts.

We validated this approach using simulated data (see Methods, Figure 1—figure supplement 3) and found that the residual variance test has high power and good control over the false positive rate when the overdispersion parameter *k* used in the null distribution was accurately estimated. To make the test more robust to factors affecting the estimation of *k* (Figure 1—figure supplement 4), we used simulation to control the false discovery rate empirically (Table 1). Our statistical test can be applied to shotgun metagenomes to sensitively and specifically identify variable genes in any environment without prior knowledge of factors that stratify relatively high versus low abundance samples.

**Table 1:**
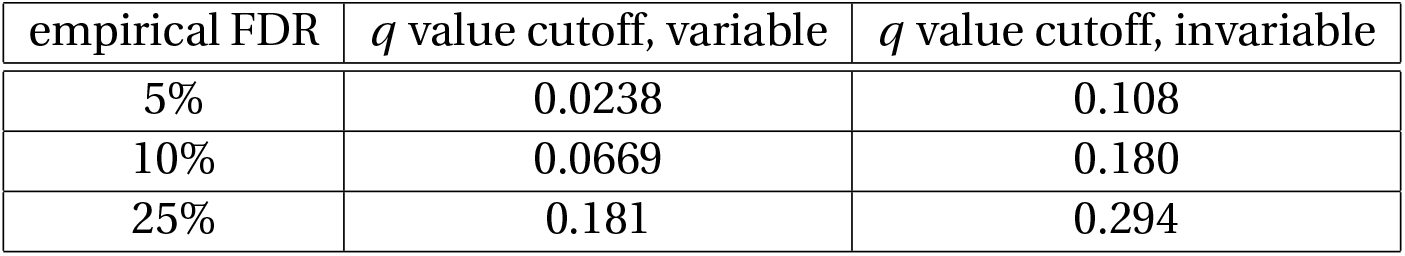
q-value cutoffs to reach a given empirical FDR, estimated from simulation.

| empirical FDR | $q$ value cutoff, variable | $q$ value cutoff, invariable |
| --- | --- | --- |
| 5% | 0.0238 | 0.108 |
| 10% | 0.0669 | 0.180 |
| 25% | 0.181 | 0.294 |

### 2.2 Thousands of variable gene families in the gut microbiome

To describe variation within healthy gut microbiota across different human populations, we randomly selected 123 metagenomes of healthy individuals from the Human Microbiome Project (HMP) [13], controls in a study of type II diabetes (T2D) [15], and controls in a study of glucose control (GC) [16]. These span American, Chinese, and European populations, respectively (see Methods). We mapped these metagenomes to KEGG Orthology families with ShotMAP [17] and counted reads for 17,417 gene families. Accurately normalizing gene read counts so that they were comparable across samples and studies is critical to our meta-analytical approach and any quantitative evaluation of shotgun metagenomes. We therefore quantified gene family abundance using log-transformed reads per kilobase of genome equivalents (log-RPKG) [18].

We found 2,357 gene families with more variability than expected and 5,432 with less (leaving 9,628 non-significant) at an empirical FDR of 5% (Figure 2—figure supplement 1). Restricting the analysis to gene families with at least one annotated representative from a bacterial or archaeal genome in KEGG, we obtained 1,219 significantly variable and 3,813 significantly invariable gene families (and 2,194 non-significant). The differences in the residual variation of these gene families can be visualized using a heatmap of the residual *ε*_*g,s*_ values (Figures 2—figure supplement 2, 2—figure supplement 3). The large number of genes that were less variable than expected given their means supports the hypothesis of some functional redundancy in the gut microbiome, potentially due to selection for core functions that make microbes more successful in the gut environment. However, our discovery of thousands of significantly variable genes across a range of abundance levels demonstrates that the gut microbiome is less invariable than prior work suggested.

This result highlights the importance of a quantitative, gene-level evaluation of functional stability. Importantly, the magnitude of the residual variance statistic 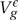 is not the sole determinant of significance, as observed by the overlap in distributions of 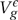 between the variable, invariable, and non-significant gene families. For example, both low-abundance gene families with many zero values and high-abundance but invariable gene families will tend to have low residual variance, but the evidence for invariability is much stronger for the second group. Our test accurately discriminates between these scenarios, tending to call the second group significantly invariable and not the first (Figure 2—figure supplement 1, inset).

### 2.3 Biological pathways contain both invariable and variable components

To test our hypothesis that the appearance of pathways and functional categories with similar abundance across samples is driven by a subset of core components, we examined individual gene variability within KEGG modules. As expected, we observed an overall signal of stability at this broad level of gene groupings. Many of the pathways previously identified as invariable (e.g., aminoacyl-tRNA metabolism, central carbon metabolism) indeed have more invariable than variable genes. However, individual genes show a much more complex picture. Even the most invariable pathways also include significantly variable genes (Figure 2). For example, the highly conserved KEGG module set “aminoacyl-tRNA biosynthesis, prokaryotes” included one variable gene at an empirical FDR of 5%, SepRS. SepRS is an O-phosphoseryl-tRNA synthetase, which is an alternative route to biosynthesis of cysteinyl-tRNA in methanogenic archaea [19]. Methanogen abundance has previously been noted to be variable between individual human guts: while DNA extraction for archaea may be less reliable than for bacteria, even optimized methods showed large standard deviations across individuals [20]. Another gene in this category was variable at a weaker level of significance (10% empirical FDR): PoxA, a variant lysyl-tRNA synthetase. Recent experimental work has shown that this protein has a diverged, novel functionality, lysinylating the elongation factor EF-P [21, 22].

**Figure 2:**
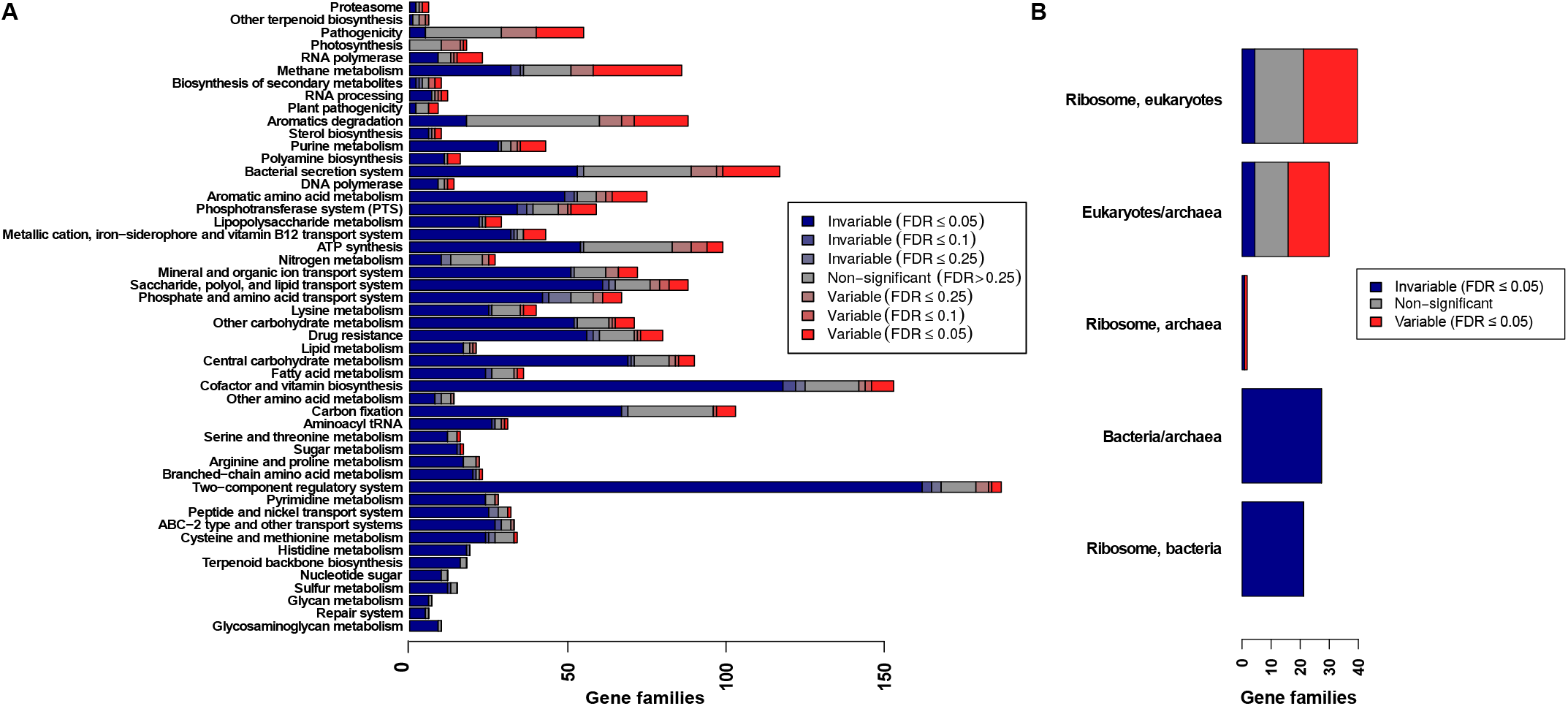
Most pathways include a mixture of both variable and invariable gene families. A) Stacked bar plots show the fraction of invariable (blue), non-significant (gray), and variable (red) gene families annotated to KEGG Orthology pathway sets (rows), at different false discovery rate (FDR) cutoffs (color intensity). Only gene families with at least one annotated bacterial or archaeal homolog are counted. B) Fraction of strongly invariable, non-significant, and strongly variable gene families within the ribosomes of different kingdoms. Row labels with only one kingdom indicate gene families unique to that kingdom, while rows with multiple kingdoms (e.g. “Eukaryotes/archaea”) indicate gene families shared between these two kingdoms. As expected, the bacterial ribosome was completely invariable.

By comparison, 77% of the tested prokaryotic gene families in the KEGG module set “central carbohydrate metabolism” were significantly invariable, and 5.6% (5 genes) were significantly variable (Figure 3—figure supplement 1) at an empirical FDR of 5%. In this case, the variable gene families highlight the complexities of microbial carbon utilization. Glucose can be metabolized by two alternative pathways: the well-known Embden-Meyerhof-Parnas (EMP) pathway (i.e., classical “glycolysis”), or the Entner-Doudoroff pathway (ED). Both take glucose to pyruvate, but with differing yields of ATP and electron carriers; ED also allows growth on sugar acids like gluconate [23]. Our analysis indicates that hosts differ in how much their gut microbial communities use ED. While all genes in the “core module” of glycolysis dealing with 3-carbon compounds were significantly invariable across individuals, we found that the ED-specific gene family edd, which takes 6-phosphogluconate to 2-keto-3-deoxy-phosphogluconate (KDPG), was significantly variable.

We also discovered significant variability in other enzymes involved in unusual sugar-phosphate and tricarboxylic acid metabolism (Figure 3—figure supplement 1). Multifunctional and primarily archaeal variants of fructose-bisphosphate aldolase (K16306, K01622) were significantly variable across hosts, while the typical FBA enzyme (FbaA) was significantly invariable. Another difference was seen in genes potentially contributing to ribose-phosphate generation. While typical pentose-phosphate pathway genes (e.g., zwf and gnd) were invariable, the bifunctional gene family Fae/Hps, thought to be involved in an alternative route to ribose-phosphate, was significantly variable [24]. Finally, a subunit of fumarate reductase, frdD, was also significantly variable. Fumarate reductase catalyzes the reverse reaction from the typical TCA cycle enzyme succinate dehydrogenase and can be used for redox balance during anaerobic growth [25]. Conversely, the standard succinate dehydrogenase genes sdhA, sdhB and sdhC were significantly invariable. These results suggest that using our test to identify variable genes within otherwise invariable pathways can reveal diverged functionality as well as families that play domain or clade-specific roles.

We found that the majority of significantly variable gene families annotated to “bacterial secretion system” (16 out of 18) were involved in specialized secretion systems, especially the type III and type VI systems (Figure 3). These secretion systems are predominantly found in Gram negative bacteria and are often involved in specialized cell-to-cell interactions, between microbes and between pathogens or symbionts and the host. They allow the injection of effector proteins, including virulence factors, directly into target cells [26, 27]. Type VI secretion systems have also been shown to be determinants of antagonistic interactions between bacteria in the gut microbiome [28, 29].

**Figure 3:**
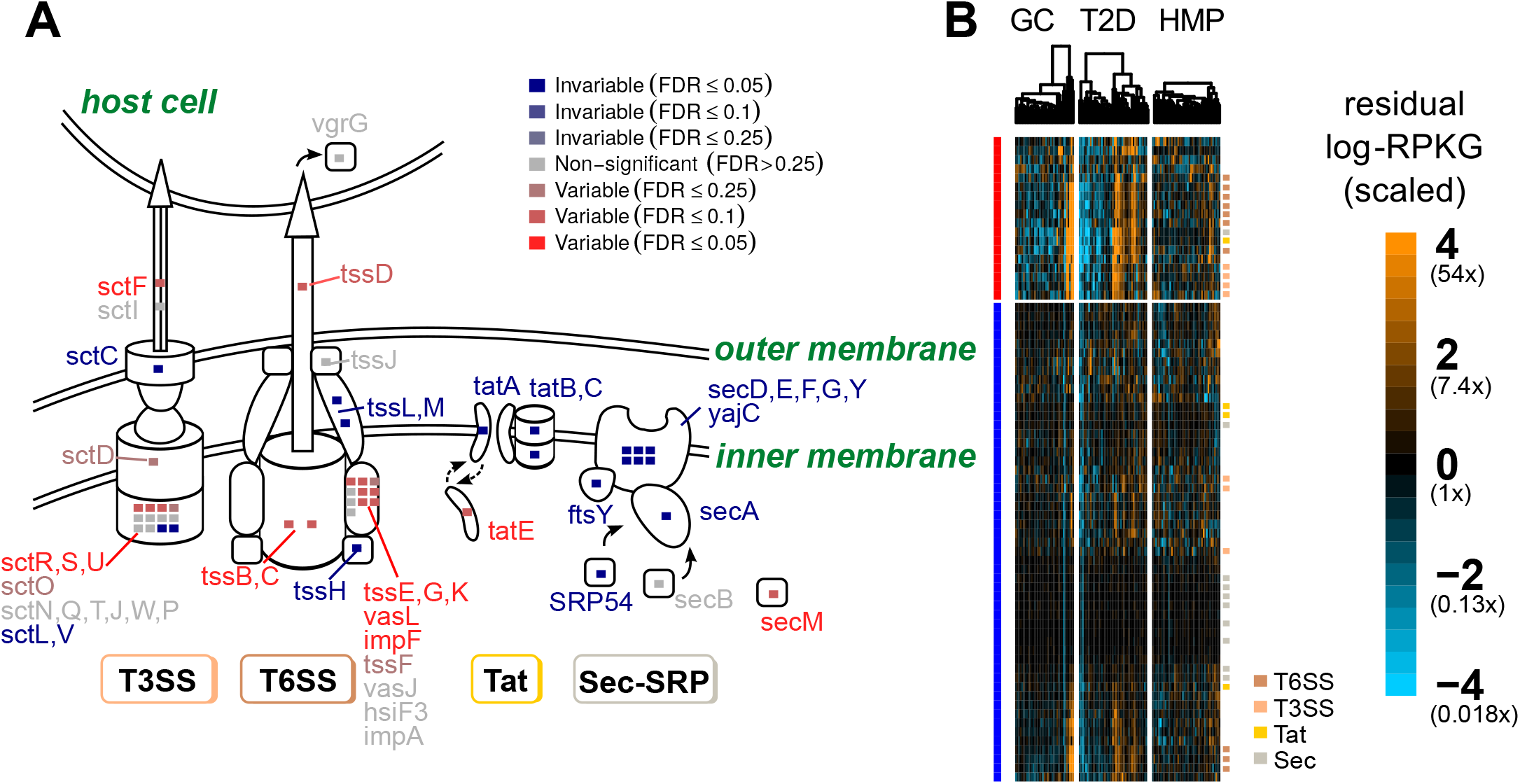
Variable and invariable gene families involved in bacterial secretion separate by gene function. A) Schematic diagram showing the type III (T3SS), type VI (T6SS), Sec, and Tat secretion system gene families measured in this dataset. Gene families are color-coded by whether they were variable (red), invariable (blue), or neither (gray), with strength of color corresponding to the FDR cutoff (color intensity). Insets show a summary of how many gene families in KEGG modules corresponding to a particular secretion system are variable or invariable and at what level of significance. B) Heatmaps showing scaled residual *log*-RPKG for gene families (rows) involved in bacterial secretion. Variable (red) and invariable (blue) gene families are clustered separately, as are samples within a particular study (columns). *log*-RPKG values are scaled by the expected variance from the negative-binomial null distribution. Genes in specific secretion systems are annotated with colored squares (T6SS: red-orange; T3SS: orange; Tat: yellow; Sec: teal).

In contrast, gene families in the Sec (general secretion) and Tat (twin-arginine translocation) pathways were nearly all significantly *invariable* at an empirical FDR of 5%, with only one gene in each being found to be significantly variable. This contradicts previous suggestions that the Sec and Tat pathways were some of the most variable in the human microbiome [13]. This discrepancy is probably due to our accounting for the mean-variance relationship in shotgun data; the Sec and Tat systems are abundant and phylogenetically diverse [30] and will therefore have high variance just by chance compared to low-abundance genes. Our test adjusts for this feature of sequencing experiments and shows that these genes are in fact less variable than expected given their mean abundance.

Our results further demonstrate that analyzing functional variability at the level of pathways can obscure gene-family-resolution trends of potential biomedical importance. The variability of individual gene families involved in lipopolysaccharide (LPS) metabolism may exemplify such a case (Figure 4). LPS (also known as “endotoxin”) is a macromolecular component of Gram-negative bacterial outer membrane, consisting of a lipid anchor called “lipid A,” a “core oligosaccharide” moiety, and a polysaccharide known as the “O-antigen” (which may be absent). Lipid A is sensed directly by the human innate immune system via the Toll-like receptor TLR4. Furthermore, lipid A variants with different covalent modifications (e.g., differentially acylated [31], phosphorylated [32], and palmitoylated [33] variants) have been shown to have different immunological properties. Hexaacylated lipid A, as found in *E. coli*, stimulates TLR4 and induces the release of pro-inflammatory cytokines; conversely, pentaacylated lipid A variants, as found in *Bacteroides*, tend not to induce TLR4 signaling, and can even prevent the hexaacylated variety from inducing inflammation [34]. This inflammation may have a variety of downstream effects on health. For example, elevated serum LPS levels are observed in obese individuals [35, 36] and individuals with inflammatory bowel disease [35], and have been linked to an increase in coronary heart disease events [37]. Conversely, a recent study advanced the hypothesis that dampening of TLR4 signaling in childhood by *Bacteroides* species may actually *increase* later susceptibility to autoimmune disease [34].

**Figure 4:**
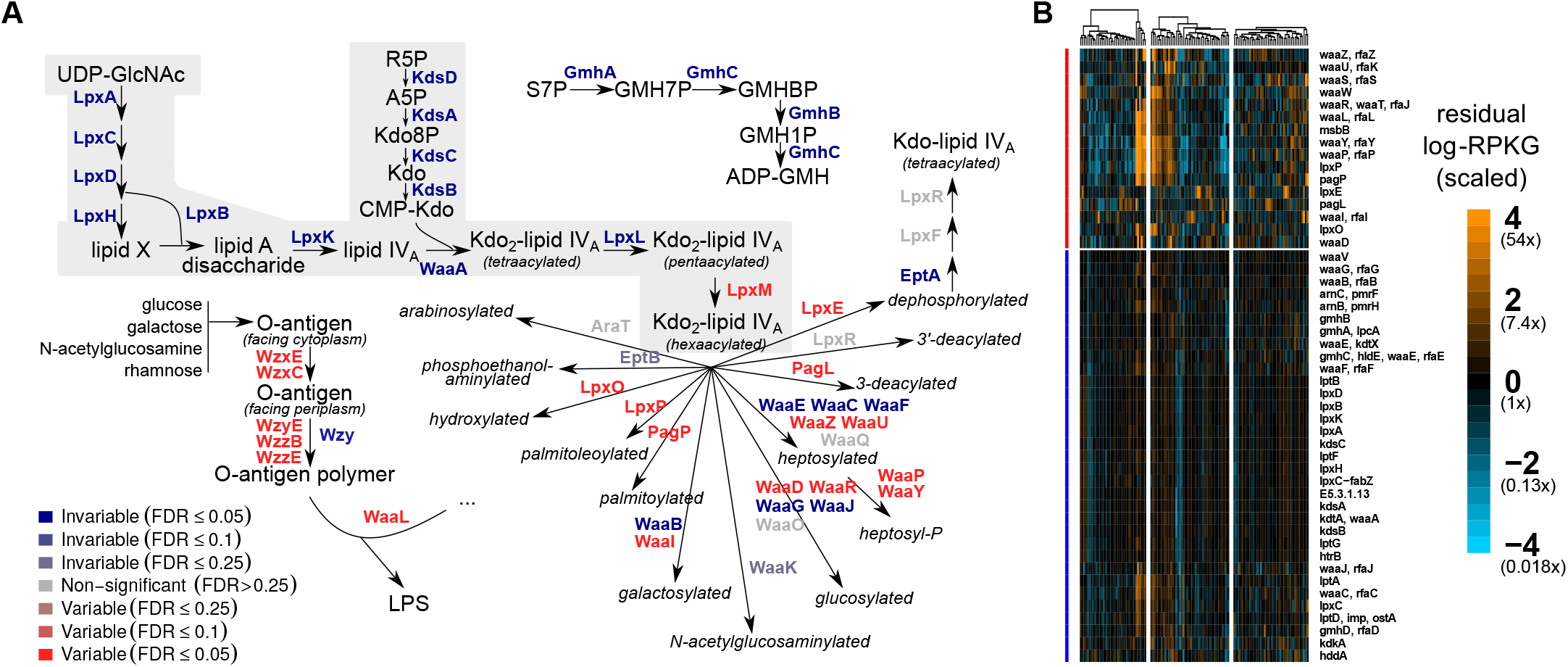
Central Kdo and lipid A biosynthesis is invariable, but many genes involved in covalent modifications to LPS are variable. A) Pathway schematic showing a selection of measured gene families involved in lipopolysaccharide metabolism. Gene families are color-coded by whether they were variable (red) or invariable (blue), with strength of color corresponding to the FDR cutoff (color intensity). Central Kdo and lipid A metabolism is highlighted in light grey. Abbreviated metabolites are GlcNAc (N-acetylglucosamine), Kdo (ketodeoxyoc-tonate), ribose-5-phosphate (R5P), sedoheptulose-7-phosphate (S7P), and glyceromannohep-tose (GMH). Aminoarabinose refers to 4-amino-4-deoxy-L-arabinose. B) Heatmaps showing scaled residual *log*-RPKG for gene families (rows) involved in lipopolysaccharide metabolism, as in Figure 3.

We found that all but one gene family involved in the biosynthesis of lipid A, as well as all gene families involved in the biosynthesis of the core oligosaccharide components ketodeoxyoctonate (Kdo) and glyceromannoheptose (GMH), were significantly invariable (16 out of 17). The lone exception catalyzes the the final lipid A acylation step, adding a sixth acyl chain; this gene family was significantly variable (FDR≤ 5%). Furthermore, we observe several variable gene families annotated as performing covalent modifications of LPS, including hydroxyl-(LpxO), palmitoyl- (PagP), and palmitoleoylation (LpxP), as well as deacylation and dephosphorylation. Previous experimental work has shown that these modifications can lead to differential TLR4 activation [33, 38]. We also observe that gene families involved in O-antigen synthesis and ligation to lipid A tended to be variable (5 out of 6). These results suggest that healthy individuals may differ in the amount of hexa- vs. pentaacylated LPS, and in the amounts of other LPS chemical modifications, and thus in their baseline level of TLR4-dependent inflammation. Importantly, since the majority of gene families annotated to LPS biosynthesis were invariable, this result would have been missed by considering the pathway as a unit.

### 2.4 Many invariable gene families are deeply conserved

Conservation of gene families across the tree of life is one factor we might expect to affect gene variability. For instance, ribosomal proteins should appear to be invariable merely because they are shared by all members of a given kingdom of life. To explore the relationship between gene family taxonomic distribution and variability in abundance across hosts, we constructed trees of the sequences in each KEGG family using ClustalOmega and FastTree. We then calculated phylogenetic distribution (PD), using tree density to correct for the overall rate of evolution [39] (Figure 5a).

**Figure 5:**
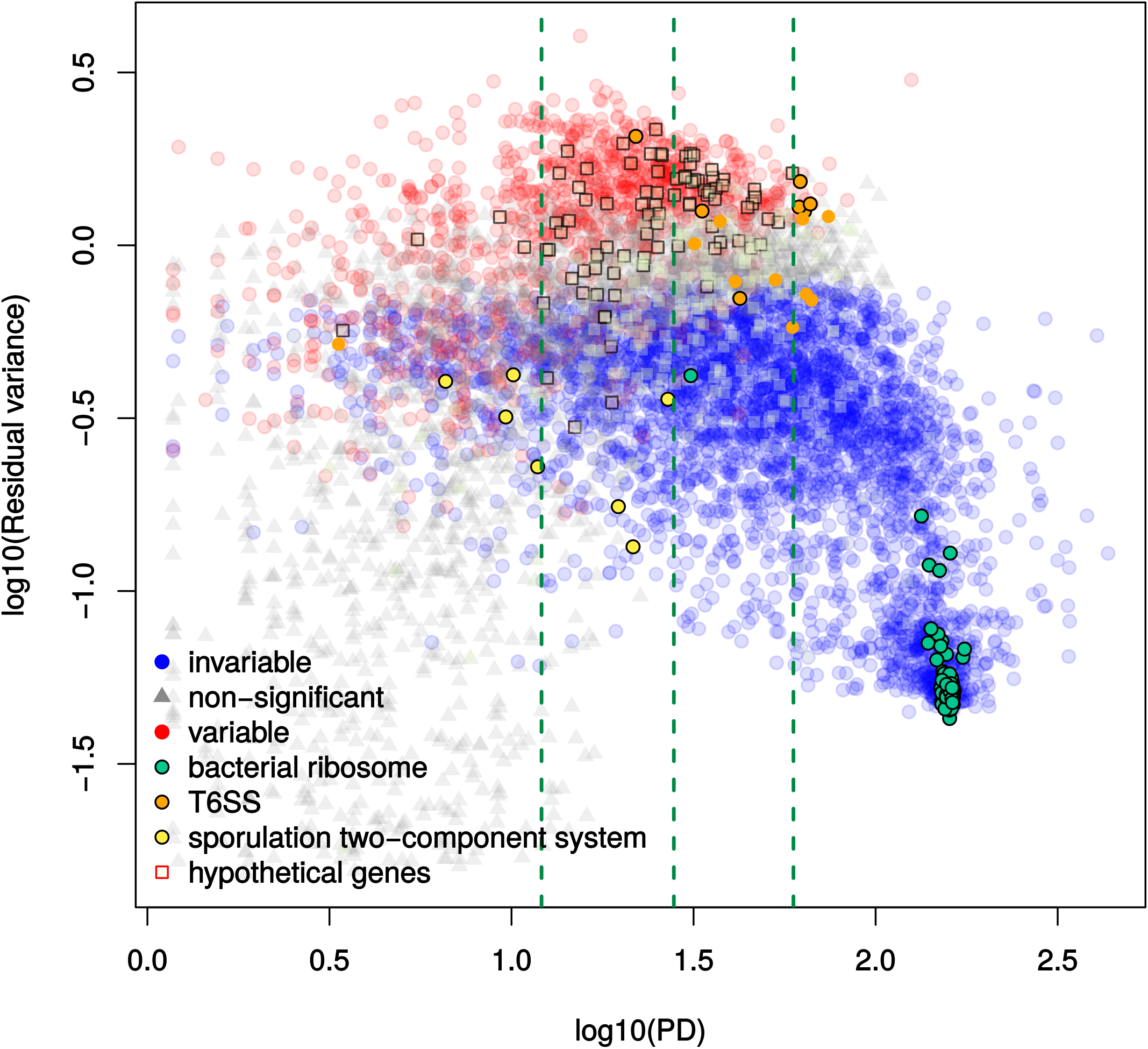
Phylogenetic distribution (PD) of gene families partially explains gene family variability. Scatter plot shows *log*_10_ PD (x-axis) vs. *log*_10_ residual variance statistic (y-axis). Red points are significantly variable while blue points are significantly invariable. Gene families in specific functional groups are also highlighted in different colors, specifically the bacterial ribo-some (green), the type VI secretion system (or “T6SS”; orange), the KinABCDE-Spo0FA sporulation control two-component signaling system (yellow), and hypothetical genes (tan squares). Gene families that were significantly invariable (ribosome and sporulation control) or significantly variable (hypothetical genes and the T6SS) at an estimated 5% FDR are outlined in black. The bacterial ribosome, as expected, had very high PD and is strongly invariable. The Type VI secretion system genes, in contrast, were conserved but variable, while some genes involved in the Kin-Spo sporulation control two-component signaling pathway have low PD but were invariable. Only gene families with at least one annotated bacterial or archaeal homolog are shown.

Overall, invariable gene families with below-median PD tended to be involved in carbohydrate metabolism and signaling. Specifically, these 2,046 gene families were enriched for the pathways “two-component signaling” (FDR-corrected p-value *q* = 1.5 × 10^−15^), “starch and sucrose metabolism” (*q =* 1.8 × 10^−3^), “amino sugar and nucleotide sugar metabolism” (*q* = 0.063), “ABC transporters” (*q* = 2.4 × 10^-5^), and “glycosaminoglycan [GAG] degradation” (*q* = 0.053), among others (Supplementary File 1). Enriched modules included a two-component system involved in sporulation control (*q* = 0.018), as well as transporters for rhamnose (*q* = 0.14), cellobiose (*q* = 0.14), and alpha- and beta-glucosides (*q* = 0.14 and *q* = 0.19, respectively). These results are consistent with the hypothesis that one function of the gut microbiome is to encode carbohydrate-utilization enzymes the host lacks [40]. Additionally, recent experiments have also shown that the major gut commensal *Bacteroides thetaiotaomicron* contains enzymes adapted to the degradation of sulfated glycans including GAGs [41, 42], and that many *Bacteroides* species can in fact use the GAG chondroitin sulfate as a sole carbon source [43].

Out of the 298 significantly-variable gene families with above-median PD, we found no pathway enrichments but three module enrichments. These included the archaeal (*q =* 1.5 × 10^-3^) and eukaryotic (*q* = 8.7 × 10^-9^) ribosomes, which reflects differences in the relative abundance of microbes from these domains of life across hosts (Figure 2b). The third conserved but variable module was the type VI secretion system (*q* = 0.039). Intriguingly, specialized secretion systems were also observed to vary within gut-microbiome-associated species in a strain-specific manner, using a wholly separate set of data [44]. Finally, gene families described as “hypothetical” were enriched in the high-PD but variable gene set (*p* = 2.4 × 10^-8^, odds ratio = 2.2) and depleted in the low-PD but invariable set (*p* = 5.4 × 10^−13^, odds ratio = 0.41).

Transporters were recently observed to show strain-specific variation in copy number across different human gut microbiomes [44], and analyses by Turnbaugh et al. identified membrane transporters as enriched in the “variable” set of functions in the microbiome [12]. However, we mainly found transporters enriched amongst gene families with similar abundance across hosts, despite being phylogenetically restricted (low-PD but invariable genes; Supplementary File 2). Part of this difference is likely due to our stratifying by phylogenetic distribution, a step previous studies did not perform.

### 2.5 Proteobacteria are the major source of variable genes

To assess which taxa contributed these variable and invariable genes, we first computed correlations between phylum relative abundances (predicted using MetaPhlAn2 [45]) and gene family abundances. This analysis revealed that the predicted abundance of Proteobacteria (and, to a lesser extent, the abundance of the archaeal phylum Euryarchaeota) tended to be correlated with variable gene families (Figure 6b).

**Figure 6:**
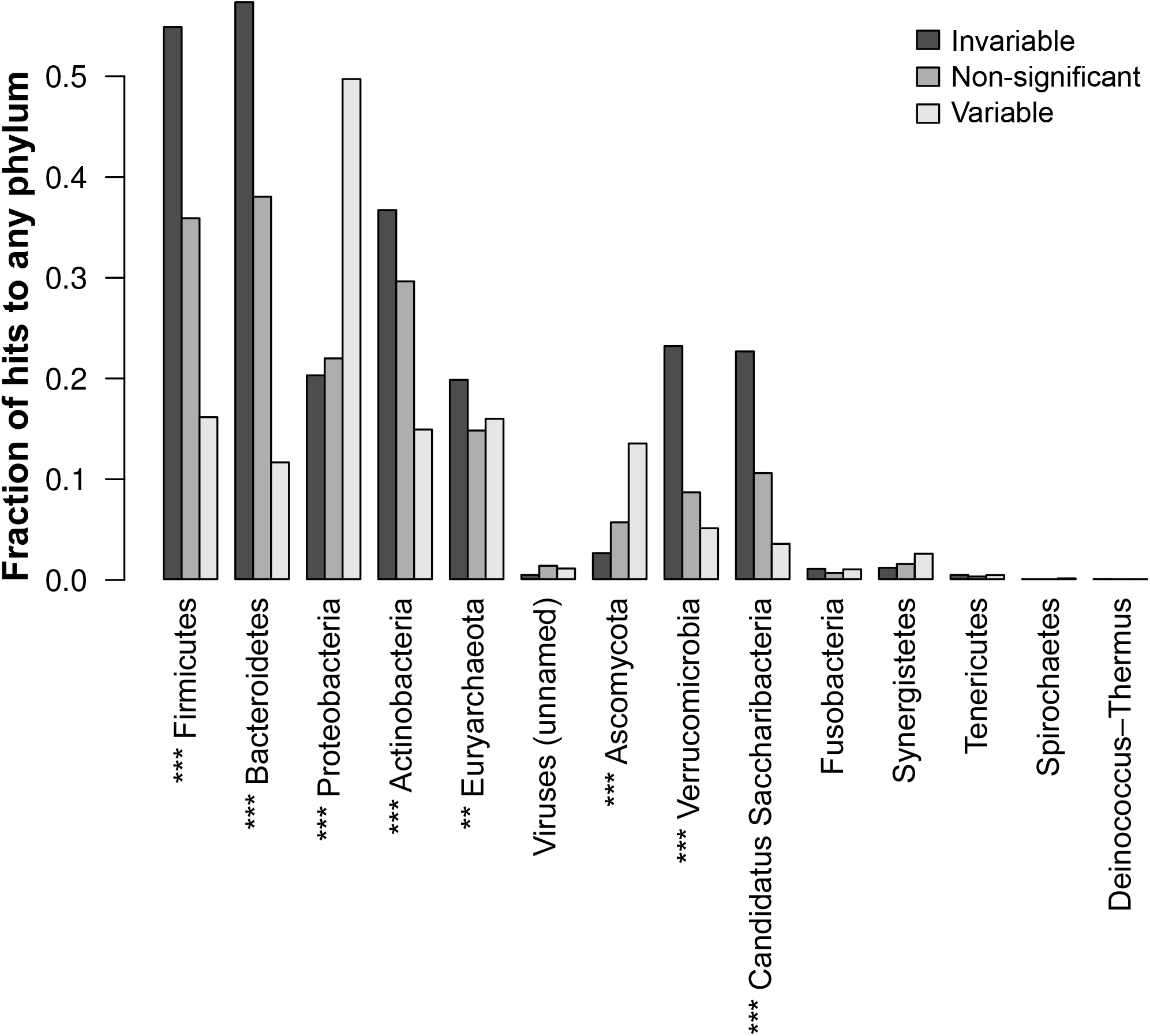
Variable gene families correlate with the predicted abundance of Proteobacte-ria. Bar plots give the fraction of gene families in each category (significantly invariable, nonsignificant, and significantly variable, 5% FDR) that were significantly correlated to predicted relative abundances of phyla, as assessed by MetaPhlan2, using partial Kendall’s τ to account for study effects and a permutation test to assess significance. Asterisks give the level of significance by chi-squared test of non-random association between gene family category and the number of significant associations. (***: *p* ≤ 10^-8^ by chi-squared test after Bonferroni correction; **: p ≤ 10^-4^.)

Proteobacteria were a comparatively minor component of these metagenomes (median = 1%), compared to Bacteroidetes (median = 59%) and Firmicutes (median = 33%). However, some hosts had up to 41% Proteobacteria. Overgrowth of Proteobacteria has been associated with metabolic syndrome [46] and inflammatory bowel disease [47]. Also, Proteobacteria can be selected (over Bacteroidetes and Firmicutes) by intestinal inflammation as tested by TLR5-knockout mice [48], and some Proteobacteria can induce colitis in this background [49], potentially leading to a feedback loop. Thus, the variable gene families we discovered could be biomarkers for dysbiosis and inflammation in otherwise healthy hosts.

We also examined correlations between gene abundance and three taxonomic summary statistics that have been previously linked to microbiome function: average genome size (AGS) [18], the Bacteroidetes/Firmicutes ratio [12, 50], and α-diversity (Shannon index). All of these statistics were *less* often correlated with variable gene families than with invariable or non-significant gene families (see Supplementary File7, Figure 6—figure supplement 1). These statistics therefore do not explain the variability of gene families in this dataset.

Finally, previous research has suggested the existence of a small number of “enterotypes” in the human gut microbiome, each with distinct taxonomic composition. A recent large-scale study confirmed that abundances of the taxa Ruminococcaceae, Bacteroides, and Prevotella explained the most taxonomic variation across individuals [51]. These enterotypes appear to be linked to long-term diet, with Prevotella highest in individuals with the most carbohydrate intake, and Bacteroides correlating with protein and animal fat. However, while these clades contribute most to taxonomic variation, all were actually *depleted* for associations with variable genes. In contrast, the Proteobacterial family Enterobacteriaceae was much more likely to be associated with variable gene families (Figure 6—figure supplement 2). This suggests that compared to previously-identified enterotype marker taxa, levels of Proteobacteria, and potentially Euryarchaeota, better explain person-to-person variation in gut microbial gene function. These less abundant phyla were missed in enterotype studies, likely because 1. enterotypes were identified by methods that will tend to weight higher-abundance taxa more, and 2. en-terotypes were identified from taxonomic, not functional data.

Because Proteobacteria are a relatively well annotated yet low abundance phylum, we explored whether either of these characteristics drive their association with variable genes. Importantly, genes correlated with Actinobacteria did not tend to be variable, even though Proteobacteria and Actinobacteria had similar levels of abundance (minimum 0%, median 1%, maximum 20%). Thus, phylum prevalence and abundance do not explain the variability of Proteobacterial genes. To investigate annotation bias, we first compared the numbers of genomes in KEGG for each phylum. There are 1,111 Proteobacterial genomes compared to 575 for Firmicutes, 276 for Actinobacteria, and only 97 for Bacteroidetes. Proteobacteria consequently had the most “private” gene families not annotated in any other phylum (1,417), compared to 538 for Firmicutes, 342 for Euryarchaeota, 215 for Actinobacteria, and 21 for Bacteroidetes. Considering only these private gene families, Proteobacteria and Euryarchaeota were enriched for variable genes, as before, whereas variable genes were depleted in the other three phyla (Figure 6—figure supplement 3). This suggests that the level of annotation does not predict the amount of variable genes. In a further test, we repeated the entire statistical test on a subset of genes, sampling one part phylum-specific genes drawn equally from Proteobacteria, Actinobacteria, Firmicutes, and Euryarchaeota, and one part genes annotated to all four phyla (see Methods). Again, Proteobacteria- and Euryarchaeota-specific genes were significantly variable more often than those from either Actinobacteria or Firmicutes (Figure 6—figure supplement 4). We therefore conclude that phylum abundance and annotation bias do not drive the enrichment of variable genes in Proteobacteria.

### 2.6 Bacterial phyla have unique sets of variable genes

The variable genefamiliesweidentifiedseemtoinclude bothgenes whosevarianceisexplained by phylum-level variation (e.g., Proteobacteria), and genes that vary within fine-grained taxonomic classifications, such as strains within species. Also, some gene families may confer adaptive advantages in the gut only within certain taxa. To detect gene families that are variable or invariable within a phylum, we repeated the test, but using only reads that mapped best to sequences from each of the four most abundant bacterial phyla (Bacteroidetes, Firmicutes, Actinobacteria, and Proteobacteria). Most (77%) gene families showed phylum-specific effects. Invariable gene families tended to agree, but the reverse was true for variable gene families: 19.4% of gene families that were invariable in one phylum were invariable in all, compared to just 0.34% (8 genes) in the variable set (Figure 7A-B). This trend was robust to the FDR cutoff (Figure 7—figure supplement 1). Gene families invariable in all four phyla were enriched for basal cellular machinery, as expected (Supplementary File 3).

**Figure 7:**
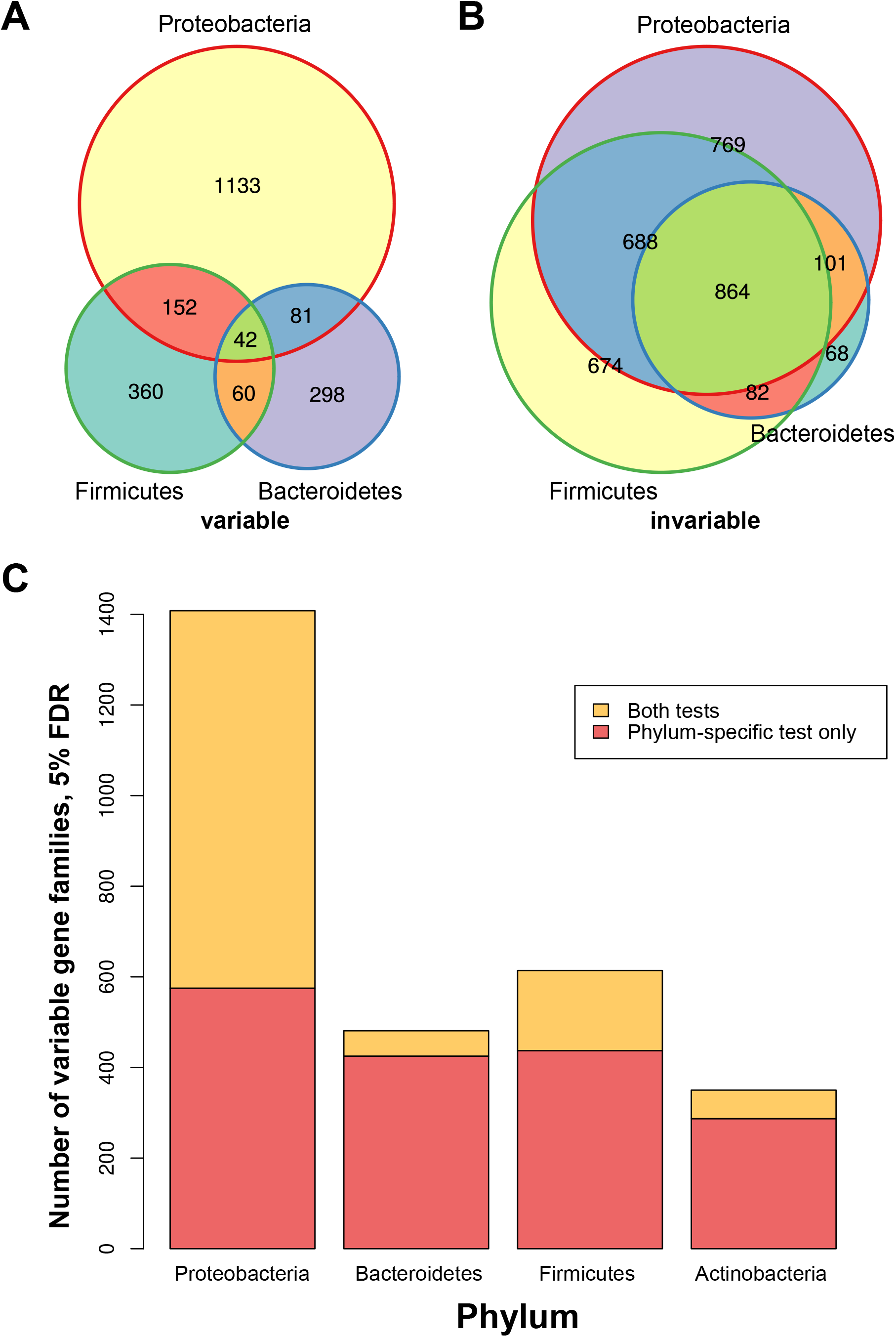
Phylum-specific tests reveal hidden variability in the most prevalent bacterial phyla. A-B) Venn diagrams showing the number of significantly variable (A) and invariable (B) gene families across Proteobacteria, Bacteroidetes, and Firmicutes, FDR ≤ 5%. C) Bars indicate the fraction of phylum-specific variable gene families that were also variable overall (yellow, “both tests”) or that were specific to a particular phylum (red, “phylum-specific test only”).

The relationship between phylum-specific and overall gene family abundance variability differed by phylum. Proteobacteria-specific variable gene families tended to be variable overall (59%), whereas the proportions of gene families that were also variable overall were much lower for Bacteroidetes- (12%), Firmicutes- (29%), and Actinobacteria-specific (18%) gene families (Figure 7C). This supports the hypothesis that Proteobacterial abundance is a dominant driver of functional variability in the human gut microbiome. It further suggests that many overall-variable gene families are not merely markers for the amount of Proteobacteria (or some other phylum), but are also variable at finer taxonomic levels, such as the species or even the strain level [44, 52].

Comparing the two dominant phyla in the gut, Bacteroidetes and Firmicutes, we further observe that the overall proportions of variable and invariable families were similar across pathways, with some interesting exceptions. For example, lipopolysaccharide (LPS) biosynthesis had many invariable gene families in Bacteroidetes and very few in Firmicutes, which we expected given that LPS is primarily made by Gram-negative bacteria. Conversely, both two-component signaling and the PTS system had many more invariable gene families in Firmicutes than in Bacteroidetes (Figure 7—figure supplement 2A). However, phylum-specific variable gene families tended not to overlap (median overlap: 0%, compared to 46% for invariable gene families). This was even true for pathways where the overall proportion of variable and invariable gene families is similar, such as cofactor and vitamin biosynthesis and central carbohydrate metabolism (Figure 7—figure supplement 2B). Thus, unique genes within invariable pathways vary in their abundance across microbiome phyla.

Furthermore, the enriched biological functions of the phylum-specific variable gene families differed by phylum (Supplementary File 4). For instance, Proteobacterial-specific variable gene families were enriched (Fisher’s test enrichment *q* = 0.13) for the biosynthesis of siderophore group nonribosomal peptides, which may reflect the importance of iron scavenging for the establishment of both pathogens (e.g. Yersinia) and commensals (E. coli) [53]. Another phylum-specific variable function appeared to be the Type IV secretion system (T4SS) within Firmicutes (*q* = 0.021). Homologs of this specialized secretion system have been shown to be involved in a wide array of biochemical interactions, including the conjugative transfer of plasmids (e.g. antibiotic-resistance cassettes) between bacteria [54]. We conclude that our approach enables the identification of substantial variation within all four major bacterial phyla in the gut, much of which is not apparent when data are analyzed at broader functional resolution or without stratifying by phylum.

### 2.7 Variable genes are not biomarkers for body mass index, sex or age

To explore associations of gene variability with measured host traits, we used a two-sided partial Kendall’s τ test that controls for study effects (Methods). Body mass index, sex, and age were measured in all three studies we analyzed. None of these variables correlated significantly with any variable gene family abundances, even at a 25% false discovery rate. This suggests that major correlates of variation in microbiota gene levels, possibly including diet and inflammation, were not measured in these studies.

## 3 Discussion

This study presents a novel statistical method that provides a finer resolution estimate of “functional redundancy” [55] in the human microbiome than was previously possible. Our test differs from previous approaches to quantifying variability in microbiome function in several key ways. First, we focus explicitly on the variability of gene family abundance, not differences in mean abundance between predefined groups, as has been done to reveal pathways whose abundance differs between body sites [56] or disease states [6]. Second, we take a finer-grained and more quantitative approach to measuring variability of microbiome functions than the studies that initially observed that biological pathways are relatively invariable [13, 12]. Our work identifies individual gene families that break this overall trend. A third important aspect of our method is that the underlying model acounts for the mean-variance relationship in count data, as well as systematic biases between studies. Finally, our null distribution is estimated from the shotgun data and does not require comparisons to sequenced genomes.

We found that basic microbial cellular machinery, such as the ribosome, tRNA-charging, and primary metabolism, were universal functional components of the microbiome, both in general and when each individual phylum was considered separately. This finding is consistent with previous results [12], and indeed, is not surprising given the broad conservation of these processes across the tree of life. In contrast, we also identified invariable gene families that have narrower phylogenetic distributions. These included, for example, proteins involved in two-component signaling, starch metabolism (including glucosides), and glycosaminoglycan metabolism. Previous experimental work has underscored the importance of some of these pathways in gut symbionts: for instance, multiple gut-associated Bacteroides species are capable of using the glycosaminoglycan chondroitin sulfate as a sole carbon source [41], and the metabolism of resistant starch in general is thought to be a critical function of the omnivorous mammalian microbiome [40]. These results suggest that the method we present is capable of identifying protein-coding gene families that contribute to fitness of symbionts within the gut. Finally, we found a number of invariable gene families whose function is not yet annotated. These gene families may represent functions that are either essential or provide advantages for life in the gut, and may therefore be particularly interesting targets for experimental follow-up (e.g., assessing whether strains in which these gene families have been knocked out in fact have slower growth rates, either in vitro or in the gut).

We also identified significantly variable gene families, including enzymes involved in carbon metabolism, specialized secretion systems such as the T6SS, and lipopolysaccharide biosynthetic genes. Proteobacteria, rather than Bacteroidetes or Firmicutes, emerge as a major source of variable genes, including some genes whose abundance also varied within the Proteobacteria (e.g., T6SS). Since Proteobacteria have been linked to inflammation and metabolic syndrome [46], we speculate that baseline inflammation may be one variable influencing functions in the gut microbiome. Some variable genes, including many of unknown function, had surprisingly broad phylogenetic distributions.

Variable gene families have a variety of ecological interpretations, e.g., first-mover effects, drift, host demography, and selection within particular gut environments. Computationally distinguishing among these possibilities is likely to present challenges. For example, distinguishing selection from random drift will probably require longitudinal data and appropriate models. Separating effects of host geography, genetics, medical history, and lifestyle will be possible only when richer phenotypic data is available from a more diverse set of human populations. To control for study bias and batch effects, it will be important to include multiple sampling sites within each study.

While statistical tests focused on differences in variances are not yet common throughout genomics, there is some recent precedent using this type of test to quantify the gene-level heterogeneity in single-cell RNA sequencing data [57, 58], and to identify variance effects in genetic association data [59]. Like Vallejos et al. [58], we model gene counts using the negative binomial distribution, and identify both significantly variable and invariable genes; in contrast, we frame our method as a frequentist hypothesis test as opposed to a Bayesian hierarchical model. Our method also accounts for study-to-study variation. Also, unlike previous approaches in this domain, the method we describe does not require biological noise to be explicitly decomposed from technical noise; our method therefore does not require the use of experimentally-spiked-in controls, which are not present in most experiments involving sequencing of the gut microbiome. Instead, we detect differences from the average level of variability using a robust nonparametric estimator, which we show through simulation leads to correct inferences under reasonable assumptions.

A similar statistical method for detecting significant (in)variability such as the one we present here could also be applied to other biomolecules measured in counts, such as metabolites, proteins, or transcripts. Performing such analyses on human microbiota would reveal patterns in the variability in the usage of particular genes, reactions, and pathways, which would expand on our investigation of potential usage based on presence in the DNA of organisms in host stool. Another important extension is to generalize our method for comparing hosts from different pre-defined groups (disease states, countries, diets) to identify gene families that are invariable in one group (e.g., healthy controls) but variable in another (e.g., patients), analogously to recent methods for the analysis of single-cell RNA-Seq [60] and GWAS [59] data. In particular, gene families whose variance differs between case and control populations could point to heterogeneity within complex diseases, interactions between the microbiome and latent variables (e.g., environmental or genetic), and/or differences in selective pressure between healthy and diseased guts. Investigating group differences in functional variability could thereby allow the detection of different trends from the more common comparison of means.

## 4 Materials and Methods

### 4.1 Data collection and processing

Stool metagenomes from healthy human guts were obtained from three sources:

1. two American cohorts from the Human Microbiome Project [13], n = 42 samples selected;
2. a Chinese cohort from a case-control study of type II diabetes (T2D) [15], n = 44 samples from controls with neither type II diabetes nor impaired glucose tolerance;
3. and a European cohort from a case-control study of glucose control [16], n = 37 samples from controls with normal glucose tolerance.

Samples were chosen to have at least 1.5 × 10^7^ reads and mode average quality scores ≥ 20 (estimated via FastQC [61]). After downloading these samples from NCBI’s Sequence Read Archive (SRA), the FASTA-formatted files were mapped to KEGG Orthology (KO) [62] protein families as previously described [17]. For consistency, each sample was rarefied to a depth of 1.5 × 10^7^ reads, and additionally, as reads from HMP were particularly variable in length, they were therefore trimmed to a uniform length of 90 bp.

For each sample, we used ShotMAP to detect how many times a particular gene family matched a read (“counts”; we added one pseudocount for reasons described below). The bit-score cutoff for matching a protein family was selected based on the average read length of each sample as recommended previously [17]. For every gene family in every sample, we also computed the average family length (AFL), or the average length of the matched genes within a family. Finally, we also computed per-sample average genome size using MicrobeCensus [18] (http://github.com/snayfach/MicrobeCensus). These quantities were used to estimate abundance values in units of RPKG, or reads per kilobase of genome equivalents [18].

These RPKG abundance values were strictly positive with a long right tail and highly correlated with the variances (Spearman’s r = 0.99). This strong mean-variance relationship is likely simply because these abundances are derived from counts that are either Poisson or negative-binomially distributed. We therefore took the natural log of the RPKG values as a variance stabilizing transformation. Because *log*(0) is infinite, we added a pseudocount before normalizing the counts and taking the log transform. Since there is no average family length (AFL) when there are no reads for a given gene family in a given sample, we imputed it in those cases using the average AFL across samples.

### 4.2 Model fitting

We fit a linear model to the data matrix of log-RPKG *D* of log-RPKG described above, with *n* gene-families by *m* samples, to capture gene-specific and dataset-specific effects:

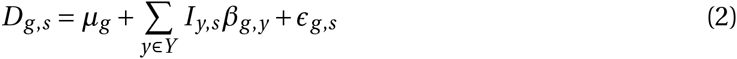

where g ∈ [1, *n*] is a particular gene family, *s* ∈ [1, *m*] is a particular sample, *µ*_*g*_ is estimated by the grande or overall mean of log-RPKG 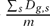 for a given gene family *g, Y* is the set of studies, *I*_*y,s*_ is an indicator variable valued 1 if sample *s* is in study *y* and 0 otherwise, *β*_*g,y*_ is a mean offset for gene family *g* in study *y*, and the residual for a given gene family and sample are given by *∈*_*g,s*_. For each gene family, the variance across samples of these *∈*_*g,s*_, which we term the “residual variance” or 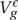, was our statistic of interest.

Overall trends in these data are explained well by this model, with an *R^2^ =* 0.20. The residuals, which are approximately symmetrically distributed around 0, represent variation in gene abundance not due to study effects.

### 4.3 Modeling residual variances under the null distribution

Having calculated this statistic 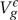 for each gene family *g*, we then needed to compare this statistic to its distribution under a null hypothesis *H*_0_. This required us to model what the data would look like if in fact there were no surprisingly variable or invariable gene families. To do this, we used the negative binomial distribution to model the original count data (before adding pseudocounts and normalization to obtain RPKG).

The negative binomial distribution is commonly used to model count data from high throughput sequencing. It can be conceptualized as a mixture of Poisson distributions with different means, which themselves follow a Gamma distribution. Like the Poisson distribution, the negative binomial distribution has an intrinsic mean-variance relationship. However, instead of a single mean-variance parameter as in the Poisson, the negative binomial can be described with two,a mean parameter and a “size” parameter, which we refer to here as *k* such that 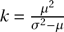. *k* ranges from (0,∞), with smaller values corresponding to more overdispersion (i.e., higher variance given the mean) and larger values approaching, in the limit, the Poisson distribution.

To model the case where no gene family has unusual variance given its mean value, i.e., our null hypothesis, we assumed that the data were negative-binomially distributed with the observed means *µ*_*g,y*_ for each gene *g* and study *y*, but where the amount of overdispersion was modeled with a single size parameter *k*_*y*_ for each study *y.* This has similarities to previous approaches to model RNAseq distributions [63] and to identify (in)variable genes from single-cell RNAseq data [58] (see also Discussion).

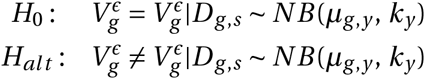

To estimate this 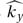, the overall size parameter for a given study *y*, we estimated the mode of per-gene-family size parameters *k*_*g,y*_ within data set *y*, using the method-of-moments estimator for each *k*_*g,y*_. We accomplished this by fitting a Gaussian kernel density estimate to the log-transformed *k*_*g,y*_ values, and then finding the 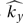 value that gave the highest density. (From simulations, we found that the mode method-of-moments was more robust than the median or harmonic mean: see Figure 1—figure supplement 2.) We could then easily generate count data under this null distribution, add a pseudocount and normalize by AFL and AGS, fit the above linear model, and obtain null residual variances 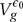 using exactly the same procedure described above.

Statistical significance was obtained by a two-tailed test:

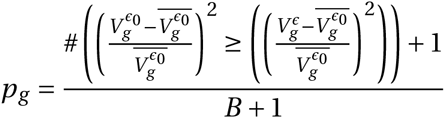

Here, *B* refers to the number of null test statistics 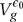 (in this case, *B* = 750), and the over-lined test statistics refer to their mean across the null distribution.

The resulting p-values were then corrected for multiple testing by converting to FDR q-values using the procedure of Storey et al. [64] as implemented in the qvalue package in R [65]. An alternative approach to determining significance is based on the bootstrap. While using a parametric null distribution allows us to explicitly model the null hypothesis, it also breaks the structure of covariance between gene families, which maybe substantial because genes are organized into operons and individual genomes within a metagenome. This structure can, optionally, be restored using a strategy outlined by Pollard and van der Laan [66]. Instead of using the test statistics 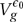 obtained under the parametric null as is, we can use these test statistics to center and scale bootstrap test statistics 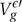, which we derive from applying a cluster bootstrap with replacement from the real data and then fitting the above linear model (2) to the resampled data to obtain bootstrap residual variances:

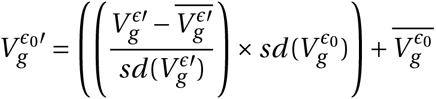

A similar non-parametric bootstrap approach has previously been successfully applied to testing for differences in gene expression [67].

### 4.4 Visualization

As expected, when the residuals are plotted in a heatmap as in Figure 2—figure supplement 2, variable gene families were generally brighter (i.e., more deviation from the mean) than invariable gene families, though not exclusively: this is because our null distribution, unlike the visualization, models the expected mean-variance relationship. We visualized this information by scaling each gene family by its expected standard deviation under the negative binomial null (i.e., by the mean root variance 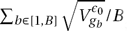) (Figure 2—figure supplement 3).

In Figure 3, for comparability with existing literature, gene families in the T6SS were named by mapping to the COG IDs used in Coulthurst [27], except when multiple KOs mapped to the same COG ID; in these cases, the original KO gene names were kept. Schematics of the T3SS, T6SS, Tat, and Sec pathways were modeled on previous reviews [68, 69, 27] and on the KEGG database [62]. The pathway diagram in Figure 4 is based on representations in the KEGG database [62], MetaCyc [70], and reviews by Wang and Quinn [71] and Whitfield and Trent [72]. These reviews were also used to identify KEGG Orthology gene families that were involved in lipopolysaccharide metabolism but not yet annotated under that term.

### 4.5 Power analysis

The test we present controls *α* as expected if the correct size parameter *k* is estimated from the data (Figure 1—figure supplement 2d-e). Estimating this parameter accurately is known to be difficult, however, particularly for highly over-dispersed data [73], and in this case we must also estimate this parameter from a mixture of true positives and nulls. We found that the mode of per-gene-family method-of-moments estimates was more robust to differences in the ratio of variable to invariable true positives (Figure 1—figure supplement 2a-c) than the median or harmonic mean (the harmonic mean mirrors the approach in Yu et al. [63]).

Power analysis was performed on simulated datasets comprising three simulated studies. For each study, 1,000 gene families were simulated over *n ∈* {60,120,480, 960} samples. Null data were drawn from a negative-binomial distribution with a randomly-selected size parameter *k* in common to all gene families, which was drawn from a log-normal distribution (log-mean= −0.65, sd= 0.57). Gene family means were also drawn from a log-normal (log mean= 2.94, sd = 2.23). True positives were drawn from a similar negative-binomial distribution, but where the size parameter was multiplied by an effect size *z* (for variable gene families) or its reciprocal 1/*z* (for invariable gene families). The above test was then applied to the simulated data, and the percent of Type I and II errors was calculated by comparing to the known gene family labels from the simulation. Using similar parameters to those estimated from our real data, we saw that α decreased and power approaches 1 with increasing sample size (see Figure 1—figure supplement 3) and that *n* = 120 appears to be sufficient to achieve control over *α*.

However, at *n* = 120, we also noted that α appeared to be greater for variable vs. invariable gene families (Figure 1—figure supplement 4), possibly because accurately detecting additional overdispersion in already-overdispersed data may be intrinsically difficult. We therefore performed additional simulations to determine *q*-value cutoffs corresponding to an empirical FDR of 5%. We calculated appropriate cutoffs based on datasets with 43% true positives and a variable:invariable gene family ratio ranging from 0.1 to 10, taking the median cutoff value across these ratios (Supplementary File 1). Using these cutoffs, the overall dataset had 45% true positives and a variable:invariable gene family ratio of 0.43.

### 4.6 Calculating phylogenetic distribution of gene families

The phylogenetic distribution (PD) of KEGG Orthology (KO) families was estimated using tree density [39]. We first obtained sequences of each full-length protein annotated to a particular KO, and then performed a multiple alignment of each family using ClustalOmega [74]. These multiple alignments were used to generate trees via FastTree [75]. For both the alignment and tree-building, we used default parameters for homologous proteins.

For all families represented in at least 5 different archaea and/or bacteria (6,703 families total), we then computed tree densities, or the sum of edge lengths divided by the mean tip height. Using tree density instead of tree height as a measure of PD corrects for the rate of evolution, which can otherwise cause very highly-conserved but slow-evolving families like the ribosome to appear to have a low PD [39]. Empirically, this measure is very similar to the number of protein sequences (Figure 5—figure supplement 1), but is not as sensitive to high or variable rates of within-species duplication: for example, families such as transposons, which exhibit high rates of duplication as well as copy-number variation between species, have a larger number of sequences than even very well-conserved proteins such as RNA polymerase, but have similar or even lower tree densities, indicating that they are not truly more broadly conserved.

Many protein families (8,931 families) did not have enough observations in order to reliably calculate tree density, with almost all of these being annotatedinonlyasingle bacterium/archaeum. For these, we predicted their PD by extrapolation. To predict PD, we used a linear model that predicted tree density based on the total number of annotations (including annotations in eukaryotes). In five-fold cross-validation, this model actually had a relatively small mean absolute percentage error (MAPE) of 13.1%. We also considered a model that took into account the taxonomic level (e.g., phylum) of the last common ancestor of all organisms in which a given protein family was annotated, but this model performed essentially identically (MAPE of 13.0%). Predicted tree densities are given in Supplementary File 6. The PD of gene families varied from 1.2 (an iron-chelate-transporting ATPase only annotated in *H. pylori*) to 434.9 (the rpoE family of RNA polymerase sigma factors).

### 4.7 Gene family enrichment

We were interested in whether particular pathways were enriched in several of the gene family sets identified in this work. For subsets of genes (such as those with specifically low PD), a 2-tailed Fisher’s exact test (i.e., hypergeometric test) was used instead to look for cases in which the overlap between a given gene set and a KEGG module or pathway was significantly larger or smaller than expected. The background set was taken to be the intersection of the set of gene families observed in the data with the set of gene families that had pathway or module-level annotations. *p*-values were converted to *q*-values as above. Finally, enrichments were enumerated by selecting all modules or pathways below *q ≤* 0.25 that had positive odds-ratios (i.e., enriched instead of depleted).

### 4.8 Associations with clinical and taxonomic variables

We were interested in using a non-parametric approach to detect association of residual RPKG with clinical and taxonomic variables (e.g., the inferred abundance of a particular phylum or other clade via MetaPhlan2). To take into account potential study effects in clinical and taxonomic variables without using a parametric modeling framework, we used partial Kendall’s *τ* correlation as implemented in the ppcor package for R [76], coding the study effects as binary nuisance variables. Kendall’s *τ* was used over Spearman’s *ρ* because of better handling of ties (an issue with taxonomic variables especially, since many, particularly at the finer-grained levels, were often zero). The null distribution was obtained by permuting the clinical/taxonomic variables within each study 250 times, and then re-assessing the partial *τ*. Finally, p-values were calculated by taking the fraction of null partial correlations equally or more extreme (i.e., distant from zero) than the real partial correlations.

Taxonomic relative abundances were predicted from the shotgun data using MetaPhlAn2 with the `--very-sensitive` flag [45].

Two approaches were used to test for annotation bias. First (Figure 6—figure supplement 3), gene families private to a phylum (i.e., those annotated in only a single bacterial/archaeal phylum) were identified from the KEGG database. We then tested whether these private gene families were enriched or depleted for significantly variable gene families (5% FDR) using Fisher’s exact test. Second (Figure 6—figure supplement 4), we performed a test in which we sampled 215 private gene families from each of Proteobacteria, Firmicutes, Actinobacteria, and Euryarchaeota, totaling 860, plus 860 gene families annotated in all four phyla. (Since Bacteroidetes only had 21 private genes, that phylum was dropped from this analysis.) Enrichment/depletion for variable gene families within each phylum was performed as above.

### 4.9 Phylum-specific tests

We created taxonomically-restricted data sets in which the abundance of each gene family was computed using only metagenomic reads aligning best to sequences from each of the four most abundant bacterial phyla (Bacteroidetes, Firmicutes, Actinobacteria, and Proteobacteria). Phylum-specific data were obtained from the overall data as follows. First, the NCBI taxonomy was parsed to obtain species annotated below each of the four major bacterial phyla (Bacteroidetes, Firmicutes, Actinobacteria, and Proteobacteria); these species were then matched with KEGG species identifiers. Next, the original RAPSearch2 [77] results were filtered, so that the only reads remaining were those for which their “best hit” in the KEGG database originally came from the genome of a species belonging to the specific phylum in question (e.g., *E. coli* for Proteobacteria). Finally, when performing the test, normalization for average genome size was accomplished by normalizing gene family counts by the median abundance of a set of 29 bacterial single-copy gene families [78], which had been filtered in the same phylum-specific way as all other gene families; this approach is similar to the MUSiCC method for average genome size correction [79]. This also controls for overall changes in phylum abundance. Finally, we estimated the average level of overdispersion 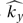 for individual studies based on the full dataset (not phylum-restricted), since the expectation that < 50% of gene families were differentially variable might not hold for each individual phylum. We used the same *q-value* cutoffs as in the overall test to set an estimated empirical FDR (Table 1). Otherwise, tests were performed as above.

### 4.10 Codebase

The scripts used to conduct the test and related analyses are available at the following URL: http://www.bitbucket.org/pbradz/variance-analyze

Counts of reads mapped to KEGG Orthology (KO) groups and average family lengths for all of the samples used in this study can be obtained at FigShare:

- https://figshare.com/s/fcflabf369155588ae41 (overall)
- https://figshare.eom/s/90d44cffdfbld214ef83 (phylum-specific)

## 5 Author contributions

PHB performed the experiments and analyses. PHB and KSP developed the test, designed the experiments, wrote the paper, and read and approved the final manuscript.

## 6 Declarations

### 6.1 Acknowledgements

The authors would like to thank Stephen Nayfach for downloading and organizing metagenomic data and metadata, and for providing and checking code for metagenome annotation, Dongying Wu for suggesting the tree density metric to measure phylogenetic distribution, Aram Avila-Herrera for help with phenotype-to-abundance associations, and Clifford Anderson-Bergman, other members of the Pollard group, and Peter Turnbaugh for helpful discussions.

### 6.2 Information about HMP clinical data

Clinical covariates for HMP were obtained from dbGaP accession #phs000228.v3.p1. Funding support for the development of NIH Human Microbiome Project - Core Microbiome Sampling Protocol A (HMP-A) was provided by the NIH Roadmap for Medical Research. Clinical data for HMP-A were jointly produced by the Baylor College of Medicine and the Washington University School of Medicine. Sequencing data for HMP-A were produced by the Baylor College of Medicine Human Genome Sequencing Center, The Broad Institute, the Genome Center at Washington University, and the J. Craig Venter Institute. These data were submitted by the EMMES Corporation, which serves as the clinical data collection site for the HMP. Authors read and agreed to abide by the Genomic Data User Code of Conduct.

Supplementary file 1: Module and pathway enrichments for variable and invariable gene sets (Fisher’s exact test *q* ≤ 0.25).

Supplementary file 2: Module and pathway enrichments for variable/high-PD and invariable/low-PD gene sets (Fisher’s exact test *q* ≤ 0.25).

Supplementary file 3: Module andpathwayenrichments for gene familieswith invariable abundances in every phylum-specific test (Fisher’s exact test, *q* ≤ 0.25).

Supplementary file 4: Module and pathway enrichments for gene families variable in each phylum-specific test (Fisher’s exact test, *q* ≤ 0.25).

Supplementary file 5: SRA IDs and characteristics (read length, average genome size from Mi-crobeCensus) for samples used in this study.

Supplementary file 6: Predicted tree densities.

Supplementary file 7: Supplementary note on correlation of variable and invariable gene families with taxonomic summary statistics

Figure 1—Source data 1: Matrix of read counts (after rarefaction) for every gene family in each sample included in the present study.

Figure 1—Source data 2: Matrix of average family lengths for every gene family in each sample included in the present study.

Figure 1—Source data 3: Log-RPKG abundances for every gene family mapped in the present study.

Figure 1—Source data 4: Residual log-RPKG abundances (i.e., after fitting the linear model) for every gene family mapped in the present study.

Figure 2—Source data 1: Counts of invariable, non-significant, and variable gene families per pathway.

Figure 2—Source data 2: Counts of invariable, non-significant, and variable gene families for ribosomes in each domain of life.

Figure 3—Source data 1: Residual log-RPKG scaled by the expected variance under the null model (see Methods).

Figure 5—Source data 1: log_10_ phylogenetic distribution (PD), log_10_ residual variance statistics (residvar), significance at 5% FDR (invariable coded as “dn”, variable coded as “up”, nonsignificant coded as “ns”), presence in at least one bacterial/archaeal genome in KEGG, and annotations for all measured gene families.

Figure 6—Source data 1: Counts of significant associations of invariable, non-significant, and variable gene families with phylum-level abundances.

Figure 6—Source data 2: Counts of significant associations of invariable, non-significant, and variable gene families with taxonomic summary statistics.

Figure 7—Source data 1: *q*-values for gene families in the overall test.

Figure 7—Source data 2: *q*-values for gene families in phylum-specific tests.

Figure 7—Source data 3: JSON-formatted lists of significantly (in)variable or non-significant gene families at 5% (“strong”), 10% (“med”), and 25% FDR (“weak”); overall test.

Figure 7—Source data 4: JSON-formatted lists of significantly (in)variable or non-significant gene families at 5% (“strong”), 10% (“med”), and 25% FDR (“weak”); phylum-specific tests.

---

**Figure 1—figure supplement 1:**
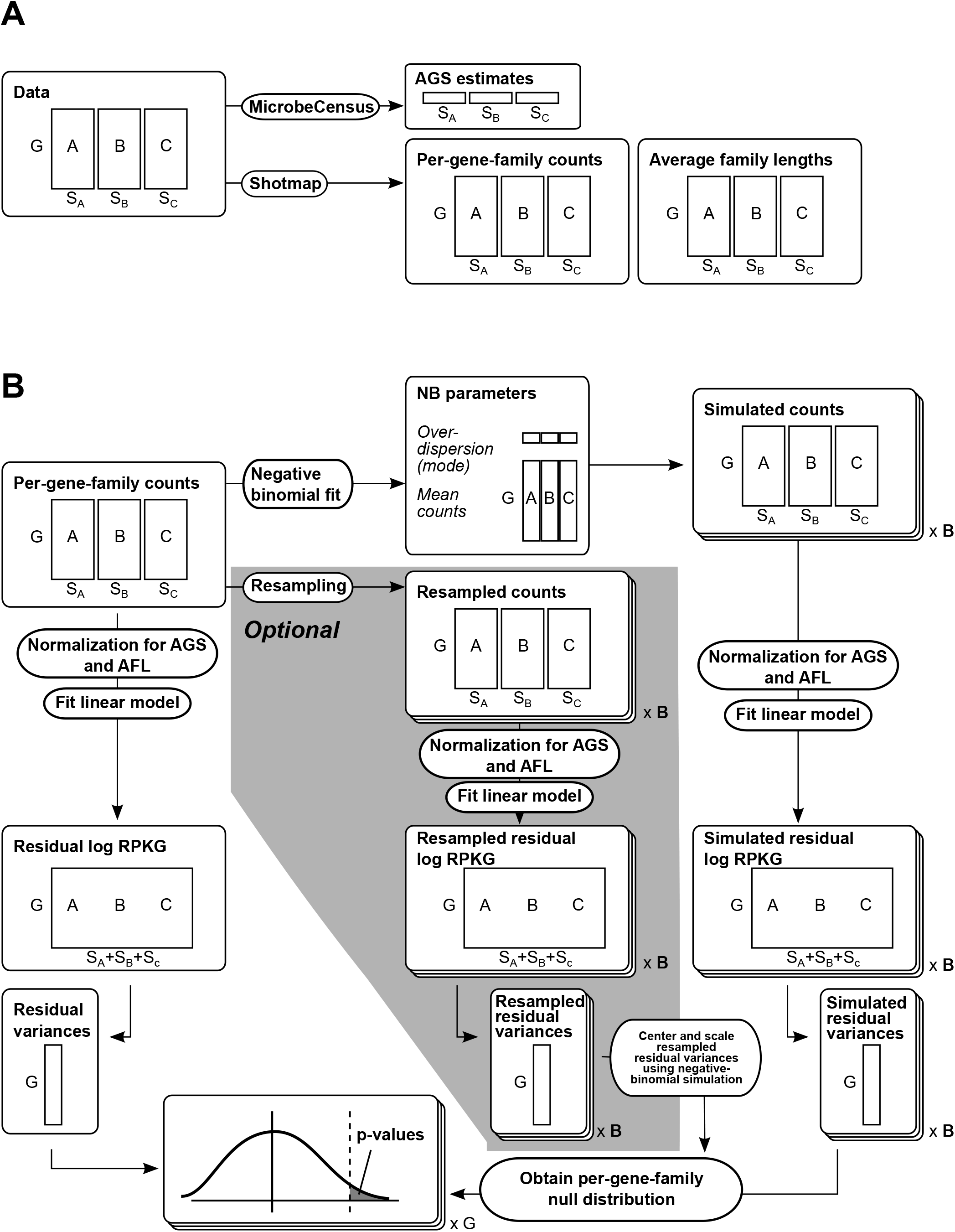
Schematic shows overview of data processing and method. A) Data is processed by taking reads from multiple datasets (represented by letters here) with a certain number of samples (represented by *S*_*A*_, *S*_*B*_, etc.). These reads will eventually map to multiple gene families G. MicrobeCensus [18] is used to estimate average genome size, while Shotmap [17] is used to map reads, yielding both matrices of counts (right hand side) and matrices of average lengths of the best-hit proteins (“average family length” or AFL). AFL and AGS estimates are used to normalize counts. B) We calculate our statistic and assign p-values as follows. First, we normalize counts from Shotmap using AFL and AGS, log-transform the resulting reads per kilobase of genome (RPKG), then applya simple linear model to fit dataset- and gene-family-specific effects. The resulting residuals (“residual log RPKG”) form a matrix of G genes by *S*_*A*_+*S*_*B*_+*S*_*C*_ samples. We take the variance across all samples for each gene to obtain a 1xG vector of residual variances. To get a null distribution, we can either use data generated fromanegative binomial fit, or, optionally, from a negative binomial fit integrated with (shaded section) bootstrap resampling. For the negative binomial fit, from the count matrices, we estimate the mean of each gene in each dataset, as well as dataset-specific overdispersion parameters *k*. We then use these to make simulated count datasets (“× B” indicating that this card is replicated once for each of B simulations), which we process as in the case of the real data, yielding simulated log-RPKG matrices and simulated residual variances for each gene family. For the resampling (if applicable), we sample with replacement from each count dataset, yielding resampled counts. We process these in the same way to obtain resampled residual variances. Finally, if using the resampled data, we center and scale the resampled residual variances using per-gene-family means and standard deviations from the simulated residual variances; otherwise, we simply take the values from applying the test to the negative binomial simulations. These form the background distribution (bottom panel, solid curve) for each gene in G (“× G” indicating that this card is replicated once for each of G genes). The actual observed residual variance (dashed line) is then compared to this distribution to obtain p-values (gray shaded area).

**Figure 1—figure supplement 2:**
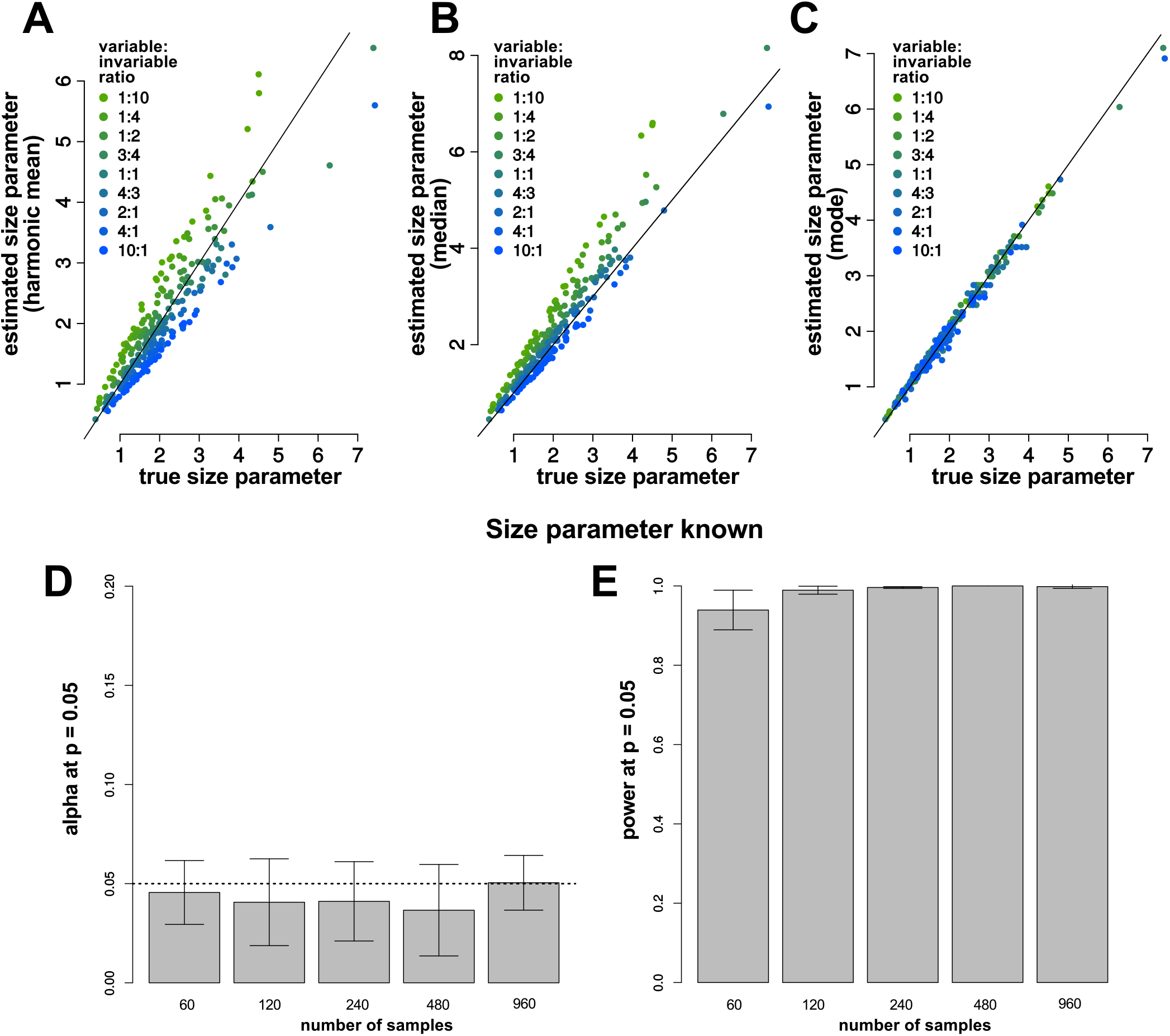
Size parameter estimator choice affects accuracy of estimation. For each mock dataset *y*, simulated null data is generated from a negative binomial distribution, fixing the size parameter *k*_*y*_ but allowing the mean *µ*_*g,y*_ to vary for each of 1,000 genes; simulated true-positive gene families are drawn from a negative-binomial distribution with size equal to *zk*_*y*_ or *k*_*y*_/*z*, where z is the effect size. A-C) The choice of estimator affects the accuracy of size estimates. The mode method-of-moments estimator (C, y-axis) more accurately estimates the true size specified in the simulation (x-axis) than the harmonic mean (A, y-axis) or median (B, y-axis), and is more tolerant to differences in the ratio of true-positive variable and invariable gene families (colors). D-E) When the size parameter is known, α (D) and power (E) are well controlled, with α approximately equal to 0.05 at p ≤ 0.05 and power approaching 1. Here, each simulation comprises three mock studies with different size parameters, mirroring our actual data. Bar heights are means from 4 simulations and error bars are ±2 SD. The proportion of variable:invariable gene families was 0.5 and 44% of genes were true positives.

**Figure 1—figure supplement 3:**
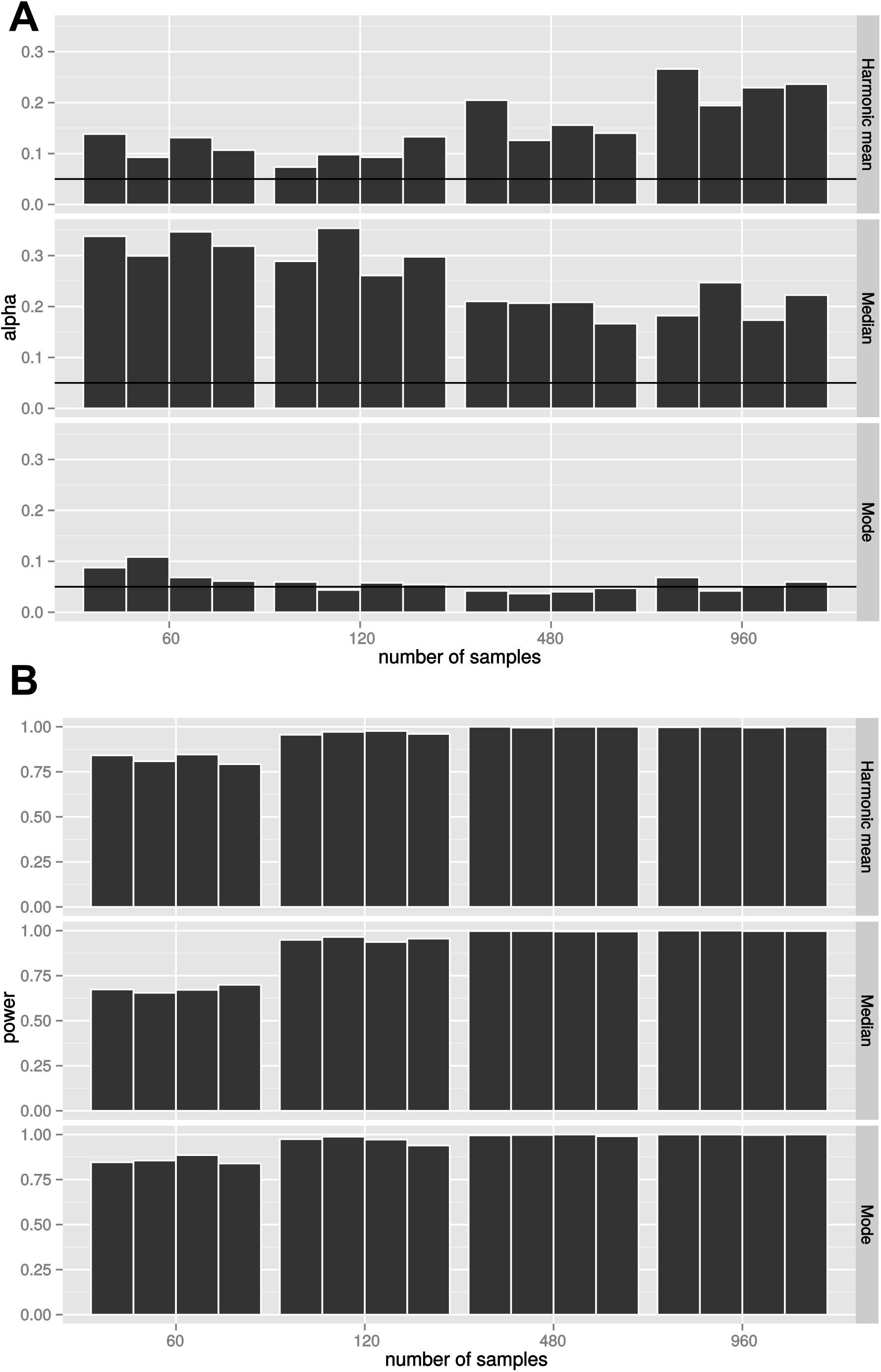
Size parameter estimation affects power and *α*, with the mode method-of-moments giving the best control. *α* (A) was minimized and power (B) was maximized when the mode method-of-moments estimator was used to get estimates of the study-specific dispersion parameters 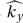. Bars are from 4 simulations. The proportion of variable invariable gene families was 0.4 and 43% of genes were true positives.

**Figure 1—figure supplement 4:**
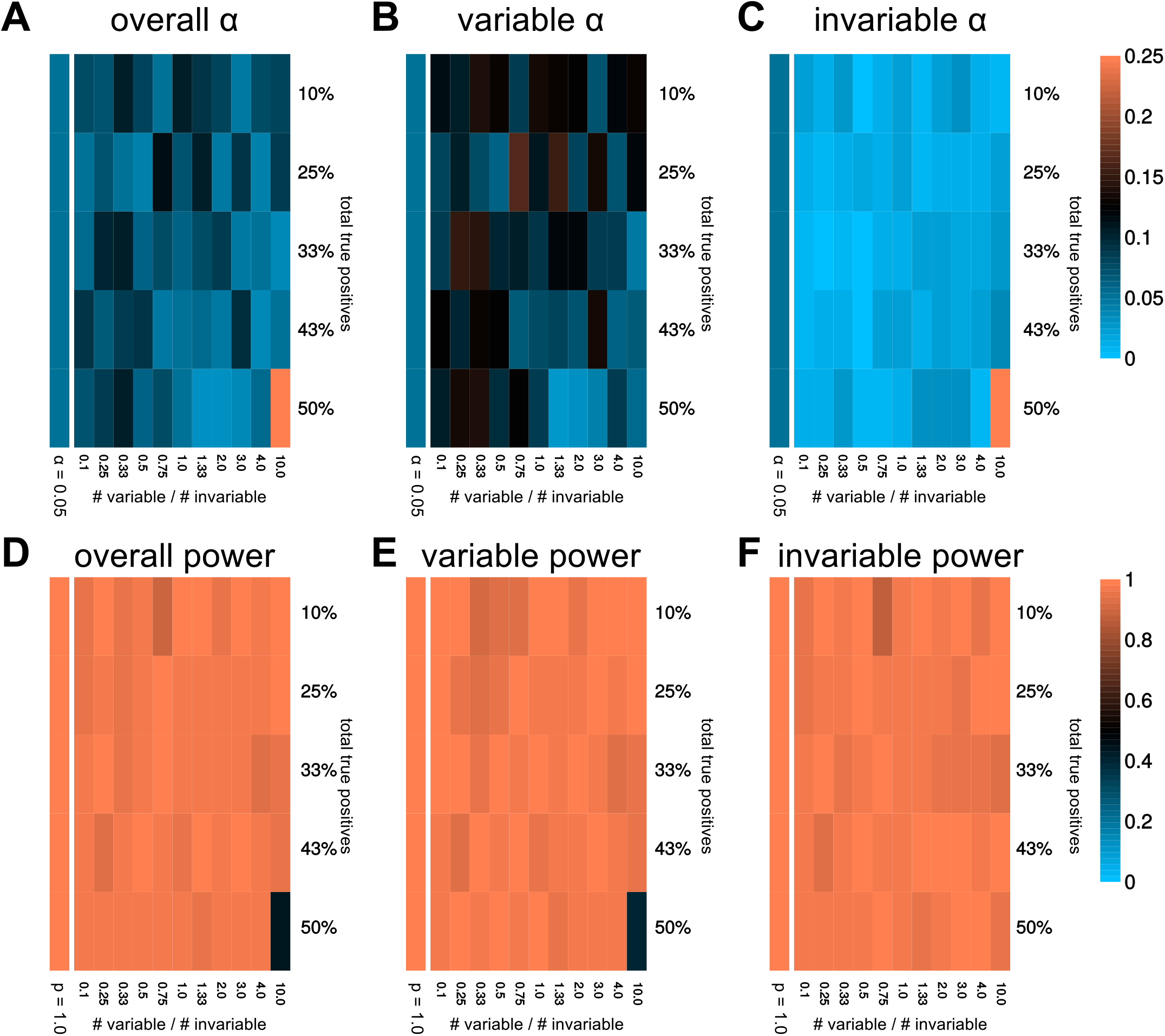
The mode estimator is robust to changes in the proportion of true positives and the ratio of variable to invariable gene families. α (A-C) and power (D-F) as a function of the proportion of true positives (x-axis) and the ratio of variable to invariable true positives (y-axis) for *n* = 120. α = 0.05 and power = 1 are shown in color-bars to the left of each heatmap for reference. α and power are calculated overall (left), for variable gene families (center), and for invariable gene families (right). In general, α was better controlled for the invariable gene families than for the variable gene families; we therefore used different empirical cutoffs for each set of genes.

**Figure 2—figure supplement 1:**
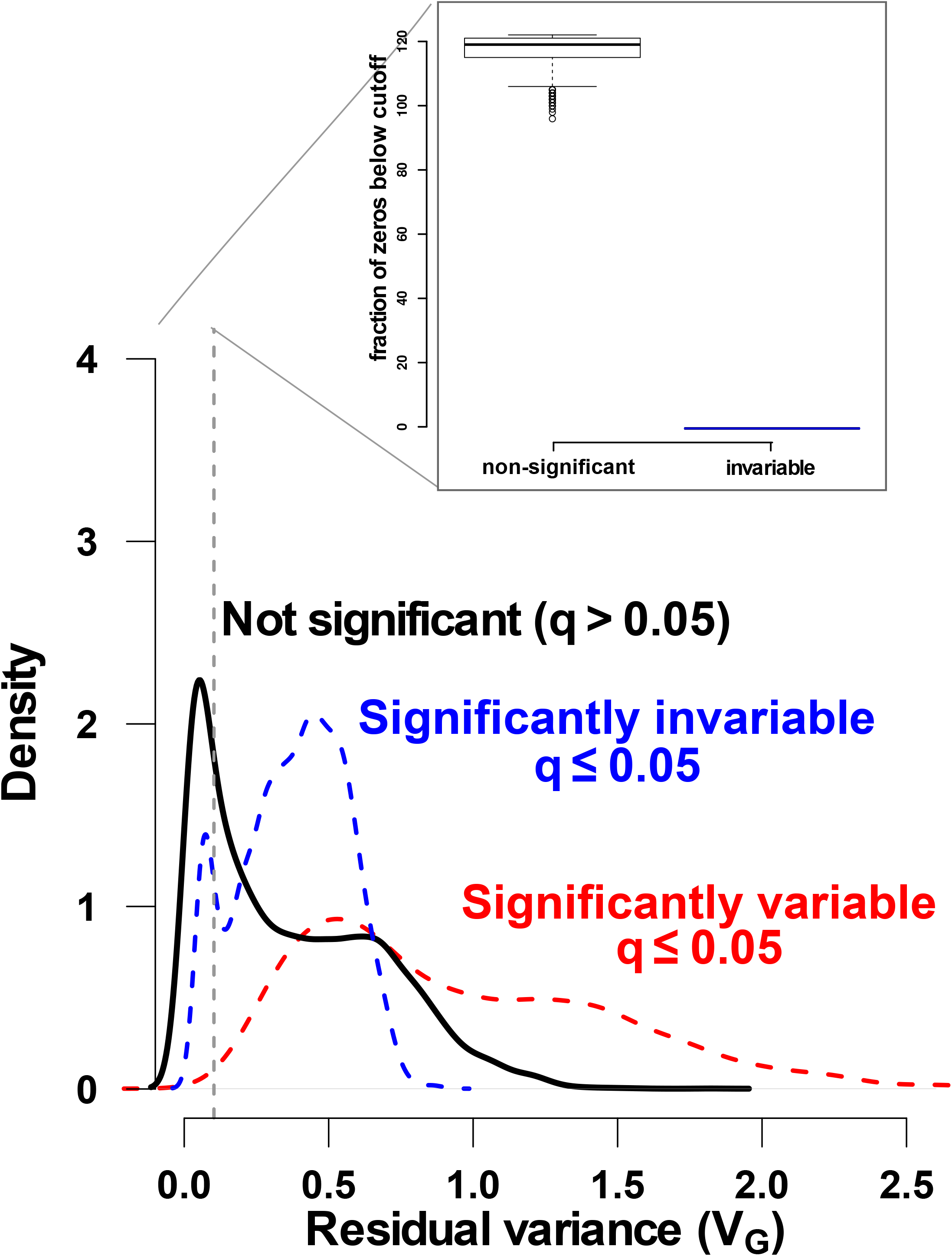
We identify significantly variable and invariable gene families. Density plots of distributions of residual variance (*V*_*G*_) statistics for significantly invariable (blue dashed line), non-significant (black solid line), and significantly variable (red dashed line) gene families. The distributions had the expected trend (e.g., significantly variable gene families tend to have higher residual variance) but also overlap, indicating the importance of the calculated null distribution. The inset shows the proportion of zero values for the non-significant (black) and significantly invariable (blue) gene families with *V*_*G*_ falling in the lowest range (vertical dashed lines), indicating that the test differentiates between gene families that only appear invariable because they have few observations, and gene families that are consistently abundant yet invariable.

**Figure 2—figure supplement 2:**
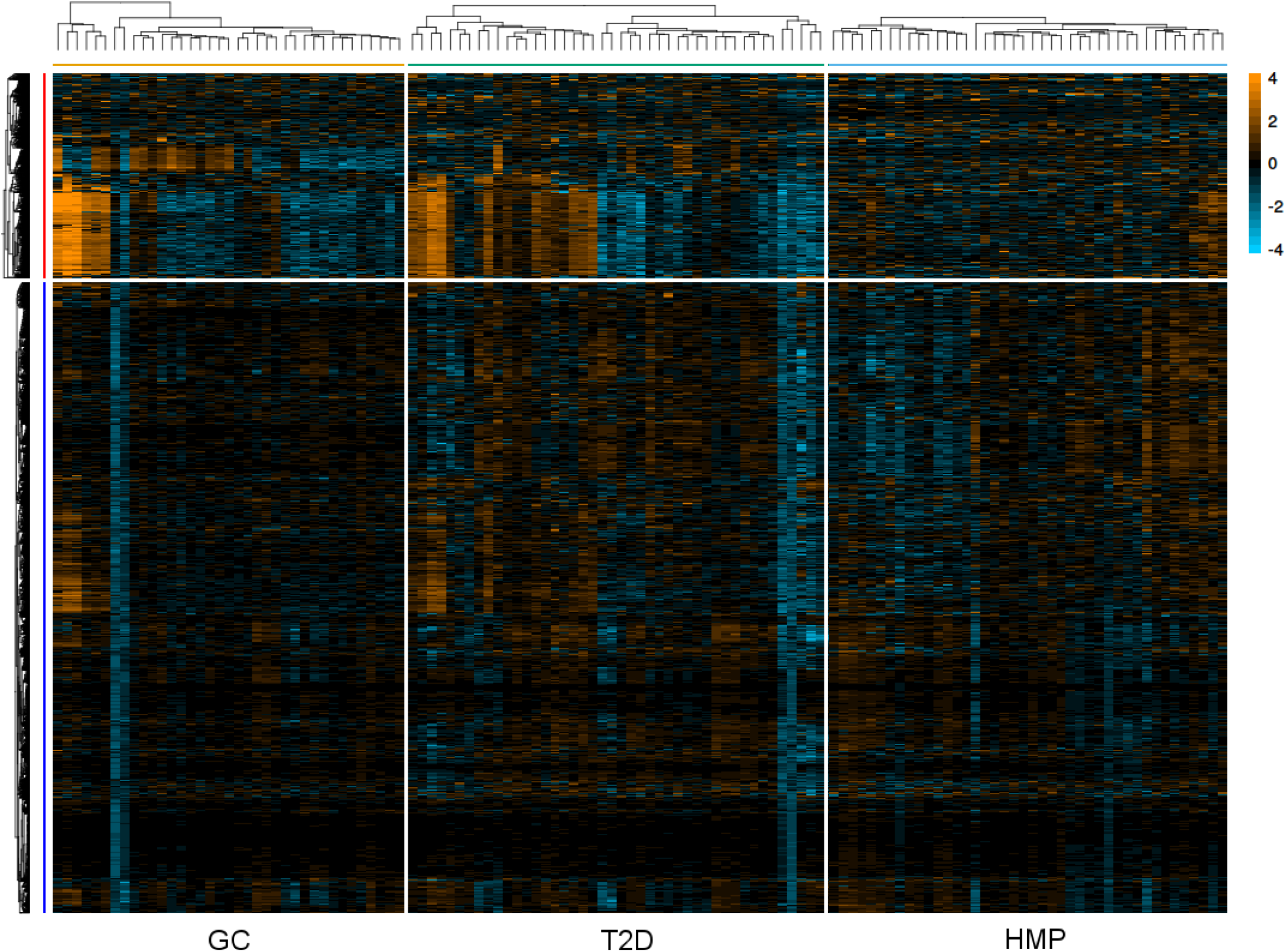
Heatmap showing significantly variable and invariable gene families (unscaled). Heatmap showing residual *log*-RPKG abundances (i.e., after normalizing for between-study effects and gene-specific abundances) of significantly invariable (blue) and significantly variable (red) gene families. Variable and invariable gene families are clustered separately, while samples are clustered within each dataset.

**Figure 2—figure supplement 3:**
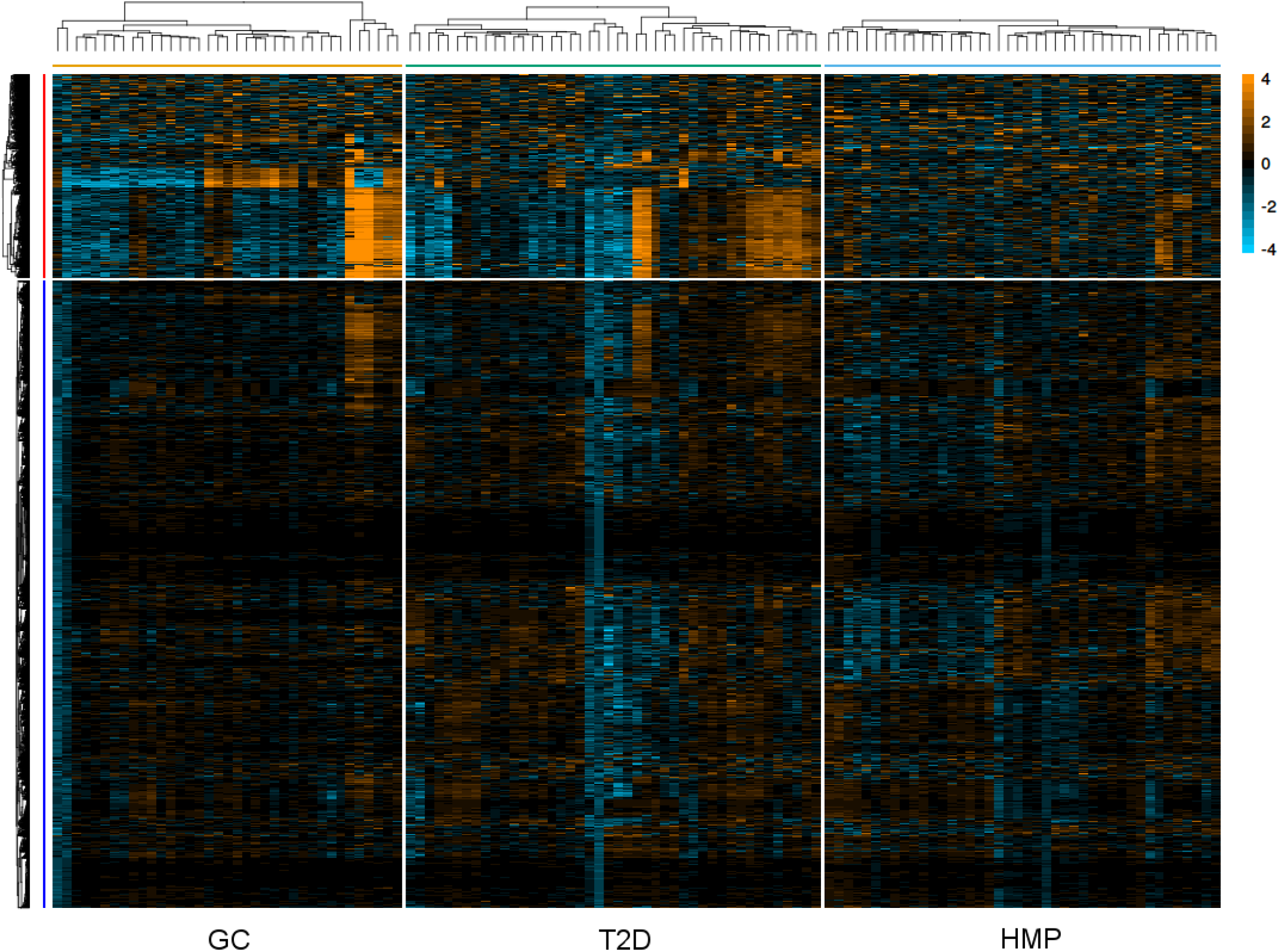
Heatmap showing significantly variable and invariable gene families (scaled). As with 2—figure supplement 2, but residual *log*-RPKG abundances scaled by their expected variance under the negative binomial null model (see Methods).

**Figure 3—figure supplement 1:**
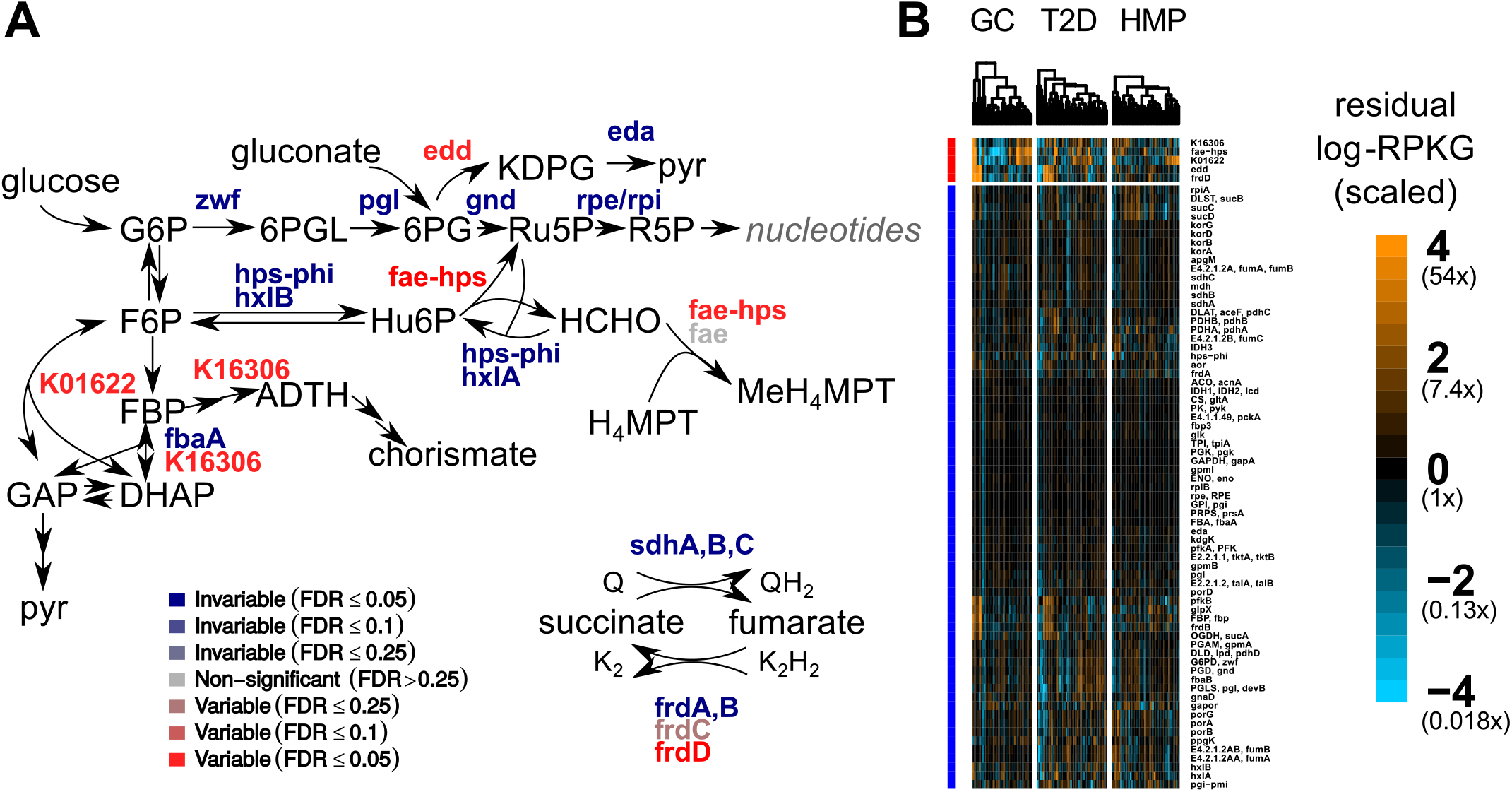
Carbon metabolism contains variable and invariable gene families. A) Pathway schematic showing a selection of measured gene families involved in central carbohydrate metabolism. Gene families are color-coded by whether they were variable (red) or invariable (blue), with strength of color corresponding to the FDR cutoff (color intensity). Genes involved in the Entner-Doudoroff pathway (edd), pentose metabolism (fae-hps), hexose metabolism (K01622, K16306), and tricarboxylic acid cycle intermediate metabolism (frdCD) were found to be variable across healthy hosts. Abbreviated metabolites are glucose-6-phosphate (G6P), fructose-6-phosphate (F6P), fructose-1,6-bisphosphate (FBP), glyceraldehyde-3-phosphate (GAP), dihydroxyacetone phosphate (DHAP), 6-phosphogluconolactone (6PGL), 6-phosphogluconate (6PG), 2-keto-3-deoxyphosphonogluconate (KDPG), ribulose-5-phosphate (R5P), ribose-5-phosphate (R5P), pyruvate (pyr), hexulose-6-phosphate (Hu6P), formaldehyde (HCHO), 2-amino-3,7-dideoxy-D-threo-hept-6-ulosonate (ADTH), and tetrahydromethanopterin (H_4_MPT). B) Heatmaps showing scaled residual *log*-RPKG for gene families (rows) involved in central carbohydrate metabolism. Variable (red) and invariable (blue) gene families are clustered separately, as are samples within a particular study (columns). *log*-RPKG values are scaled by the expected variance from the negative-binomial null distribution.

**Figure 5—figure supplement 1:**
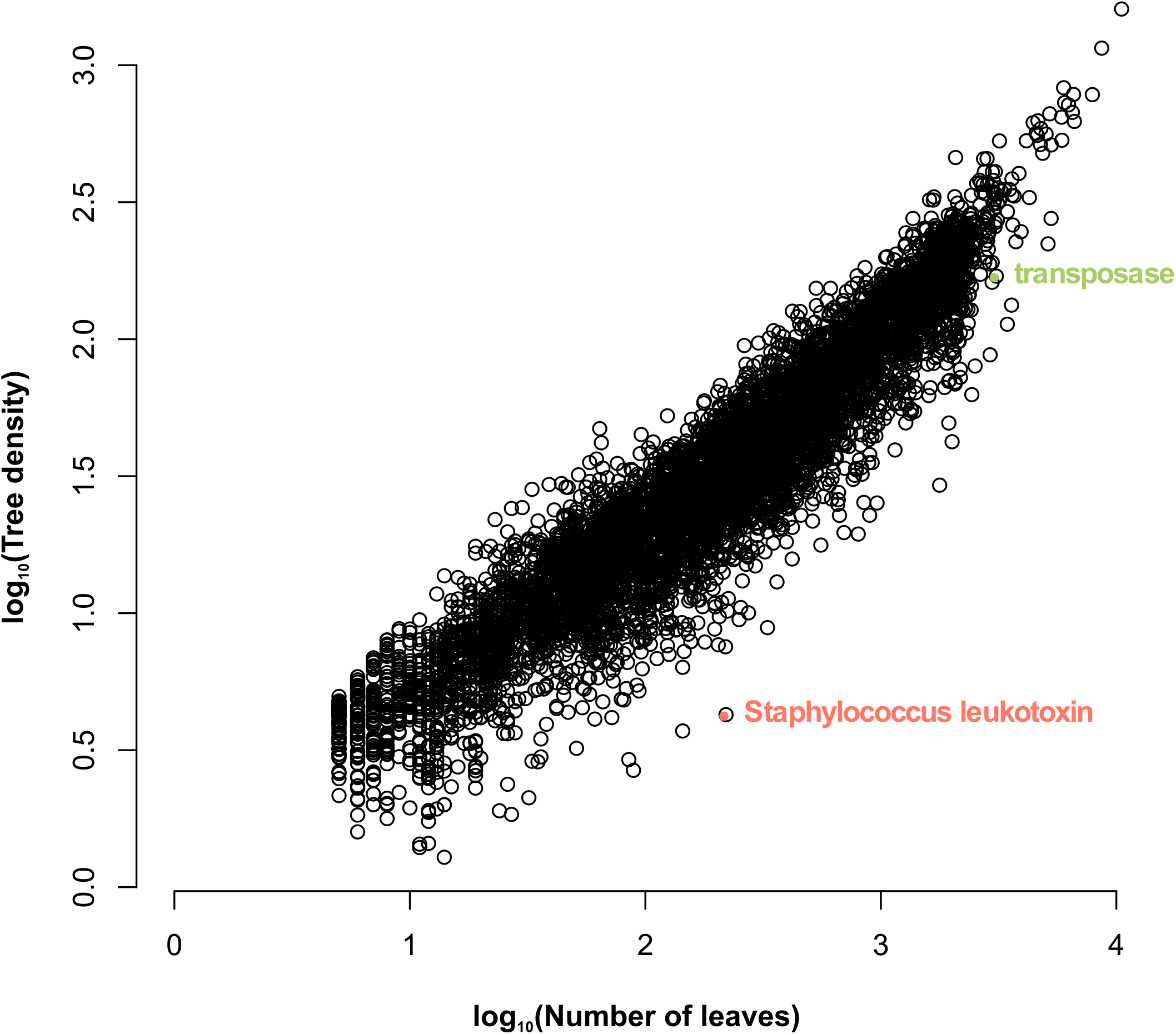
Number of leaves is correlated with tree density, but tree density corrects for the overall rate of evolution. The number of leaves (i.e., individual sequences) is plotted vs. tree density on a log-log scatter plot, with each circle representing one gene family. Two outliers with lower density than expected are plotted in colors: a putative transposase (green) and a *Staphylococcus* leukotoxin (red). Both families have large numbers of sequences from the same organism.

**Figure 6—figure supplement 1:**
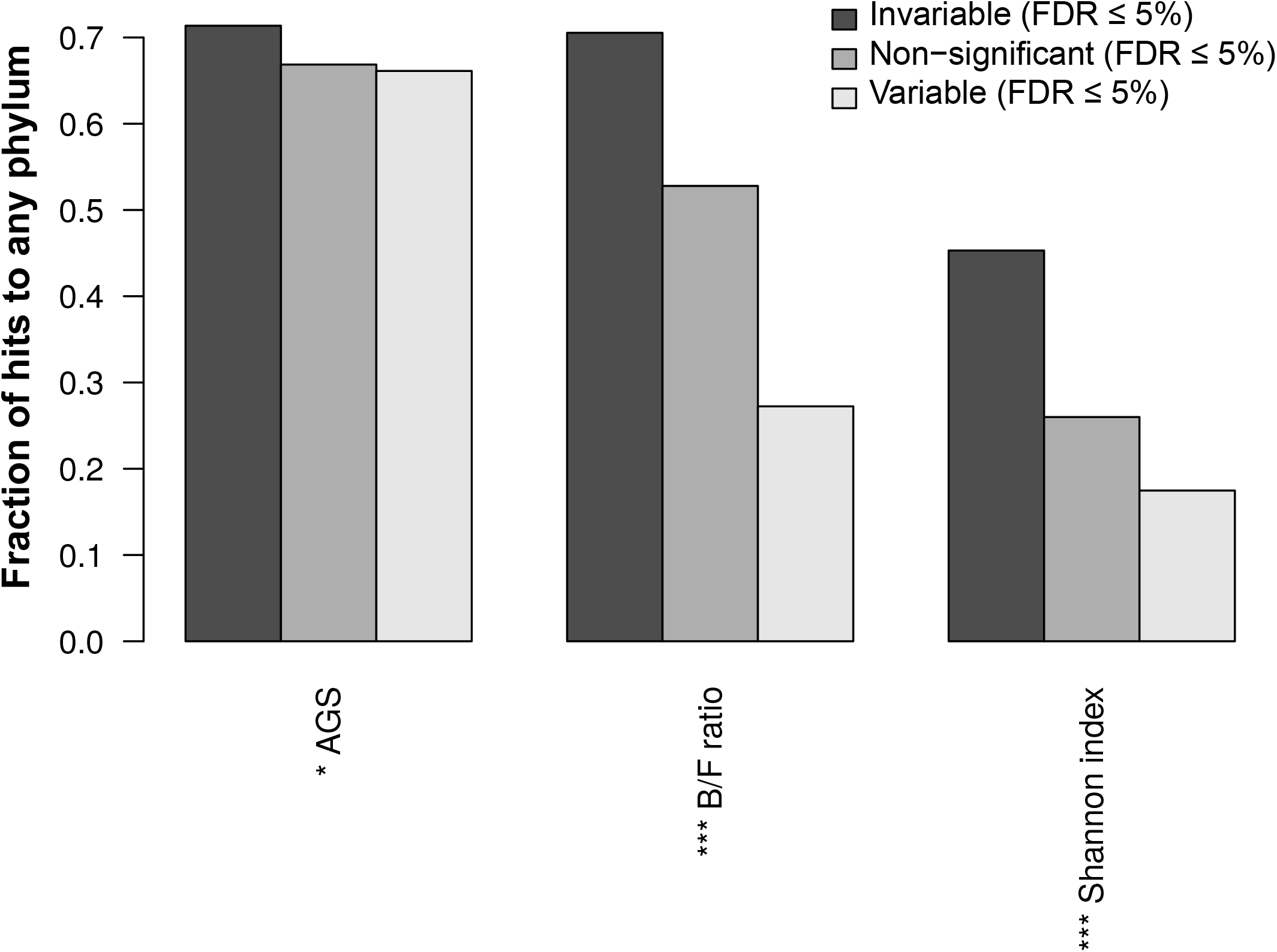
Variable gene families are less-often correlated to measured host characteristics. Bar plots give the fraction of gene families with at least one bacterial or archaeal representative in each category (significantly invariable, non-significant, and significantly variable) that were significantly correlated to various sample characteristics, using partial Kendall’s τ to account for study effects and a permutation test to assess significance. These sample characteristics are average genome size (AGS), the ratio of Bacteroidetes to Firmicutes (B/F ratio), and a measure of α-diversity (Shannon index). (***: p ≤ 10^-8^ by chi-squared test after Bonferroni correction; **: p ≤ 10^-4^.)

**Figure 6—figure supplement 2:**
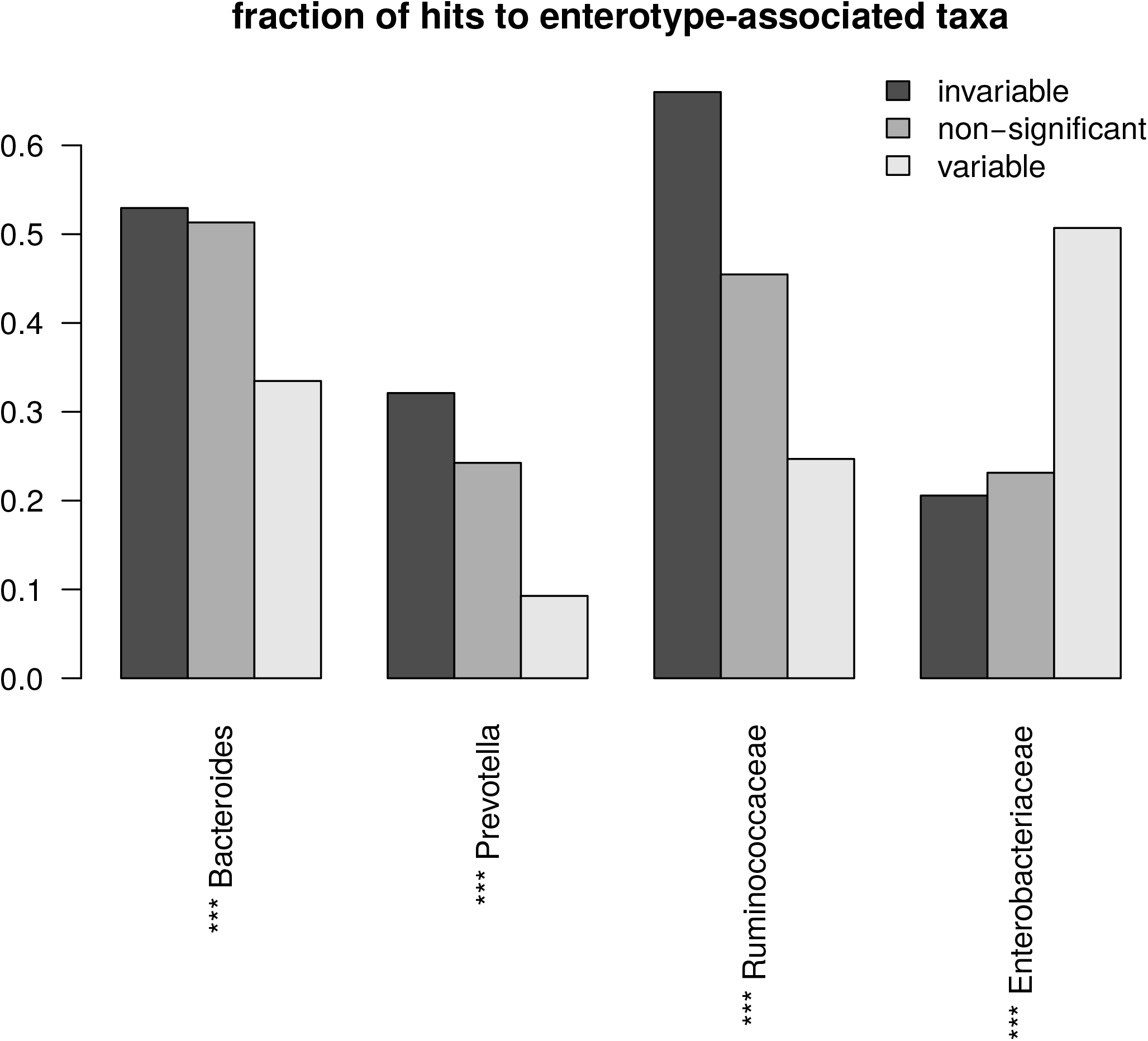
Variable gene families are less often correlated to enterotype-associated taxa, and more often correlated to the Proteobacterial clade *Enterobacteriaceae*. Bar plots give the fraction of gene families with at least one bacterial or archaeal representative in each category (significantly invariable, non-significant, and significantly variable) that were significantly correlated to the predicted abundance of specific bacterial clades (the genera Bacteroides and Prevotella, and the families Ruminococcaceae and Enterobacteriaceae). Significance was assessed as in Figure 6—figure supplement 1. (***: p ≤ 10^-8^ by chi-squared test after Bonferroni correction; **: p ≤ 10^-4^.)

**Figure 6—figure supplement 3:**
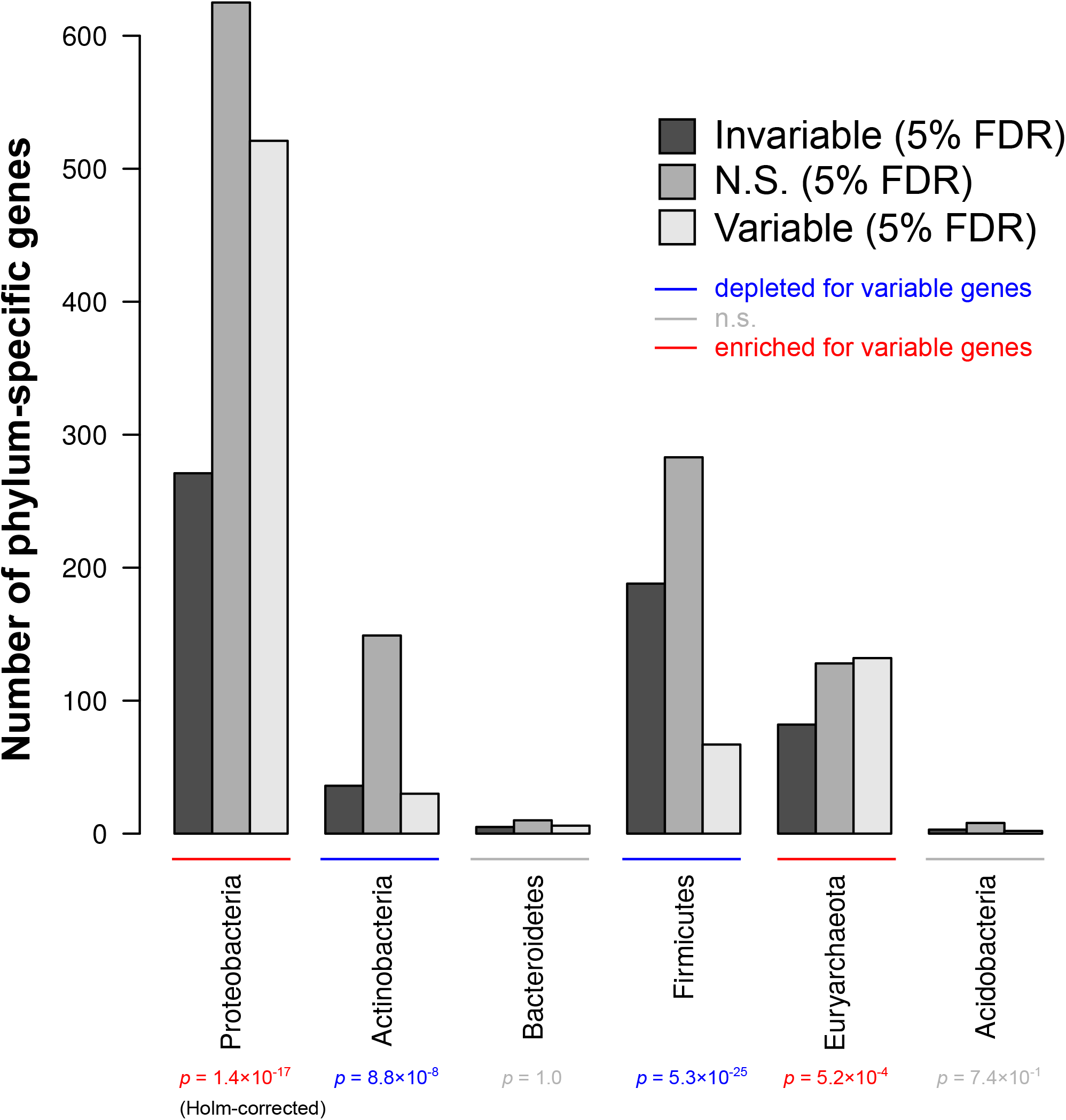
Genes only annotated in Proteobacteria or Euryarchaeota, but not Actinobacteria or Firmicutes, are more likely to be variable. Bar plots give the fraction of gene families with at least one bacterial or archaeal representative in each category (significantly invariable, non-significant, and significantly variable) that were annotated *only* in the phylum listed (x-axis). Significance was assessed as in Figure 6—figure supplement 1, using a Holm correction for significance. p-values are color-coded by whether a phylum was enriched (red), depleted (blue), or neither (gray) for variable gene families (Holm-corrected p ≤ 0.1).

**Figure 6—figure supplement 4:**
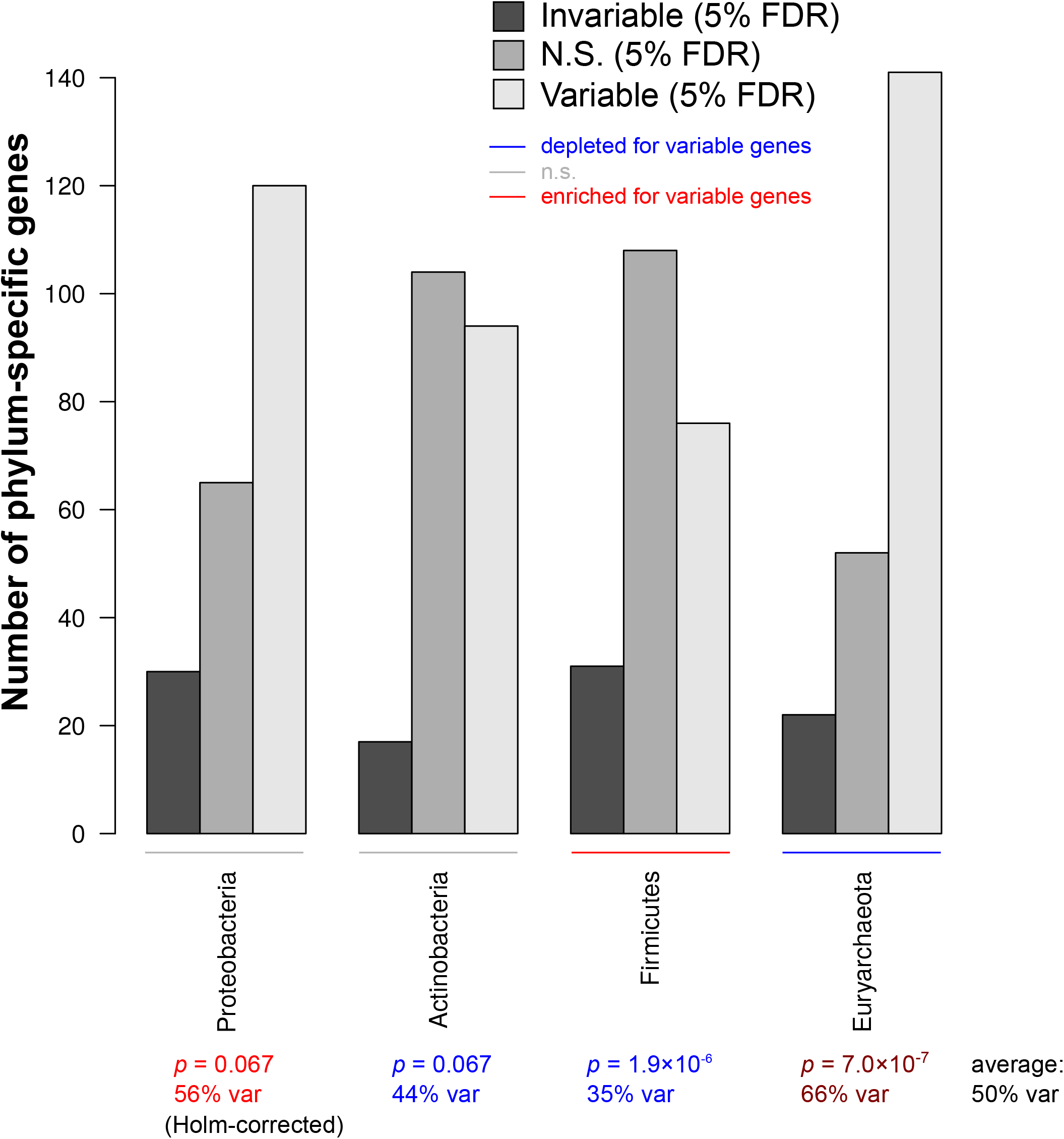
Genes only annotated in Proteobacteria or Euryarchaeota, but not Actinobacteria or Firmicutes, are more likely to be variable in a test that uniformly samples from phylum-specific genes. Bar plots are as per Figure 6—figure supplement 3, but test results come from a test that sampled equal parts phylum-specific genes and genes annotated in all four listed phyla, with phylum-specific genes themselves uniformly sampled across phyla. Significance was assessed as in Figure 6—figure supplement 3. p-values are color-coded by whether a phylum was enriched (red), depleted (blue), or neither (gray) for variable gene families (Holm-corrected *p* ≤ 0.1).

**Figure 7—figure supplement 1:**
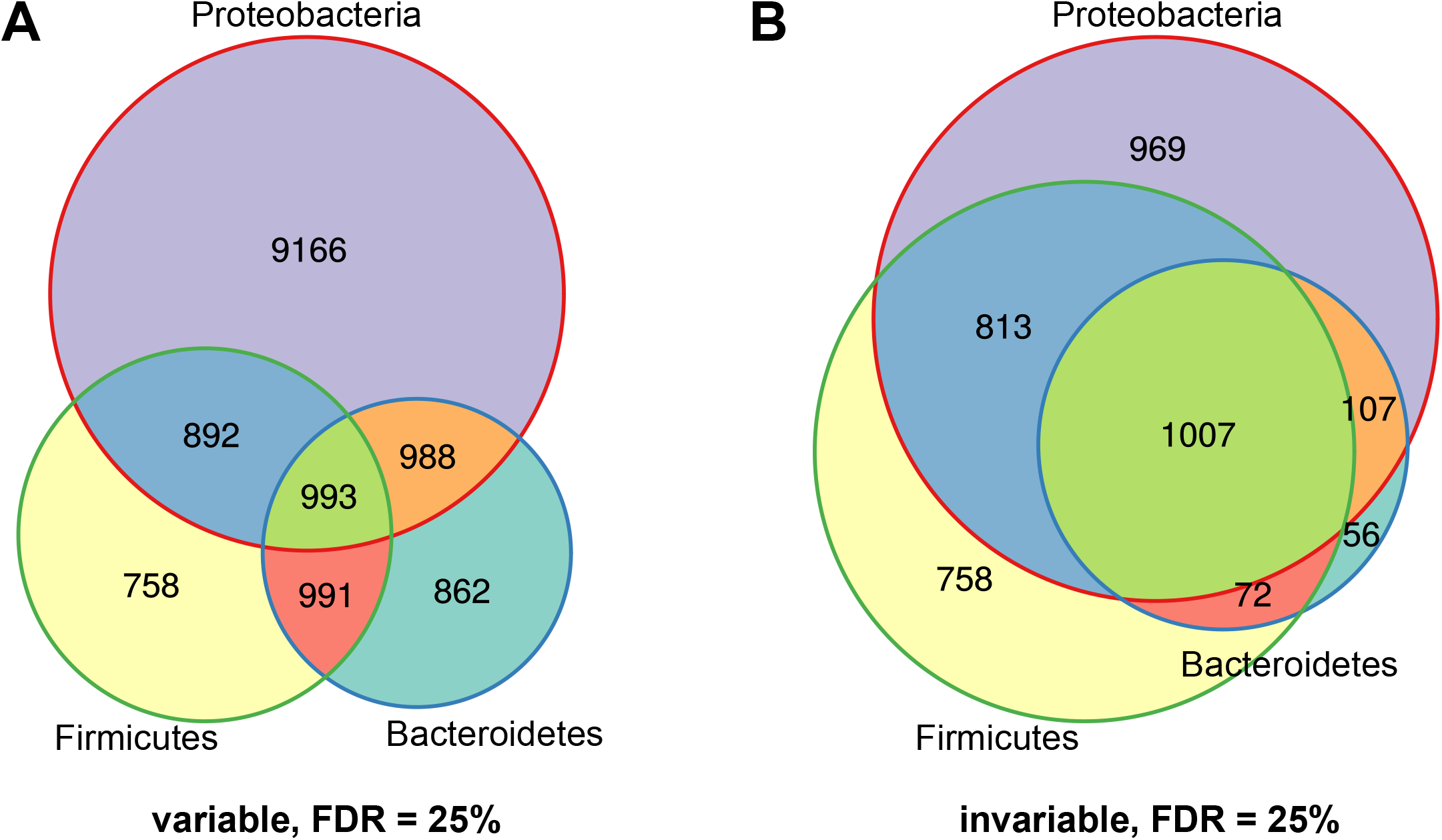
Phyla show similar trendsofoverlapatagenerous FDR cutoff. A-B) Venn diagrams showing the number of significantly variable (A) and invariable (B) gene families across Proteobacteria, Bacteroidetes, and Firmicutes, FDR ≤ 25%. Compare to Figure 7A-B.

**Figure 7—figure supplement 2:**
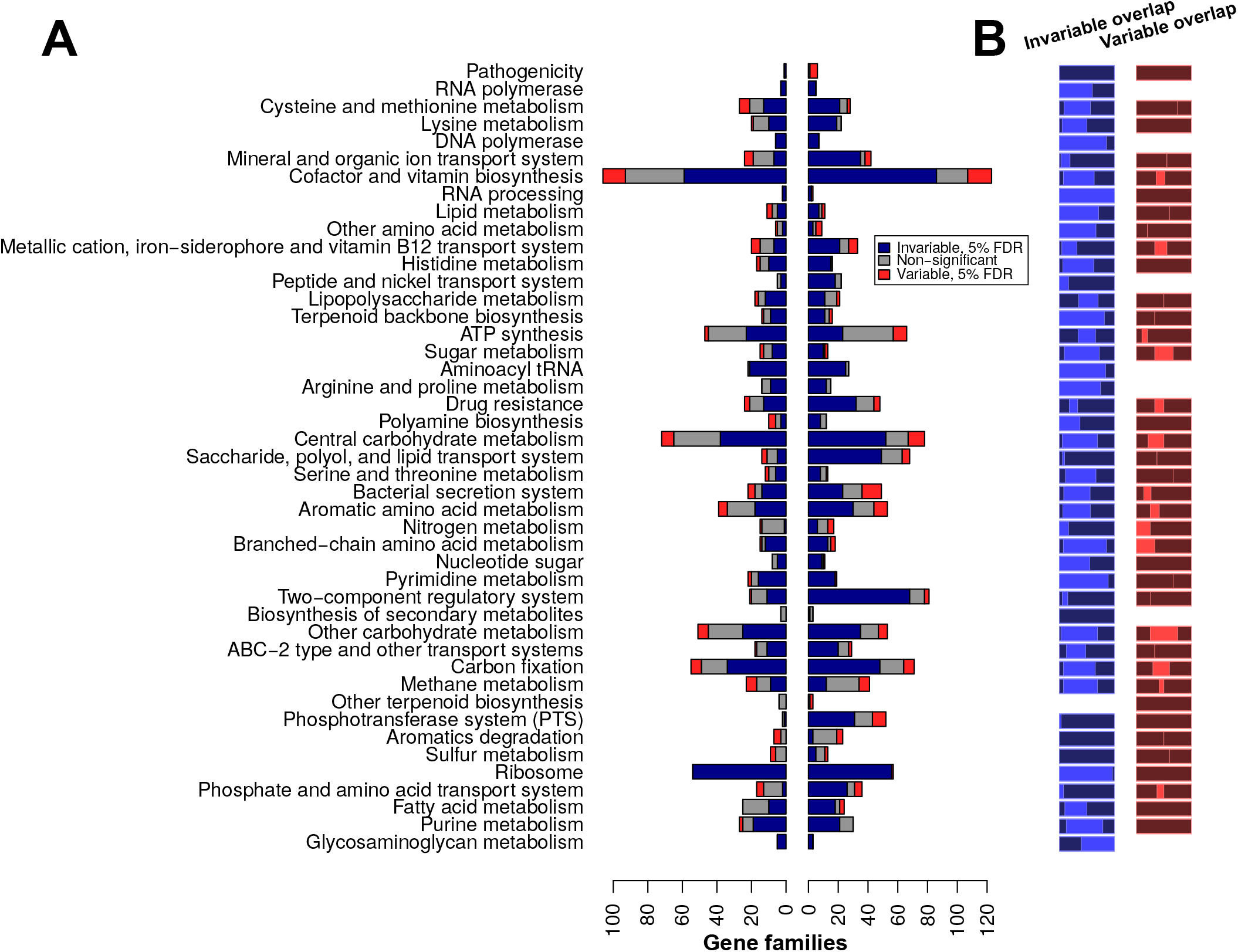
Comparison between Bacteroidetes- and Firmicutes-specific variable and invariable genes. A) Bars indicate the fraction of phylum-specific variable gene families that were also variable overall (red, “both tests”) or that were specific to a particular phylum (yellow, “phylum-specific test only”). A) For the Bacteroidetes- (left) and Firmicutes-(right) specific tests, the proportion of invariable (blue), non-significant (gray), and variable (red) gene families, at an estimated 5% FDR (using cutoffs from overall test). Pathways with at least 5 total gene families across both phyla are shown. B) Rectangular Venn diagrams showing the proportion of Bacteroides-specific (left), shared (center, bright), and Firmicutes-specific (right) invariable (blue) and variable (red) gene families for each of the pathways enumerated in B.

## Supplemental Information

### 1 Correlation of variable and invariable gene families with taxonomic summary statistics

It has previously been suggested [1] that the genome size of gut microbiota reflects a trade-off between specialization (in which metabolic pathways for the production of reliably present nutrients may be lost over time, potentially resulting in auxotrophy) and generalization, or the ability to survive and grow in different metabolic conditions (which may require more biosynthetic genes). AGS itself has also been linked to health outcomes; for instance, individuals with Crohn’s disease tend to have gut microbiota with larger genome size [2]. However, variable gene families were no more likely to be associated with AGS. Only 66% of variable gene families (with at least one bacterial or archaeal representative) had abundances that were significantly correlated with average genome size (*q ≤* 0.05), compared to 71% of invariable gene families and 66% of non-significant families at the same threshold. Thus, genome size correlates generally with gene abundance but does not predict variability of genes in healthy hosts.

The most dominant phylum-level trend across healthy human gut microbiomes is the trade-off between the two dominant phyla, Bacteroidetes and Firmicutes. The ratio of these two phyla (B/F ratio) has been linked to obesity in some studies [3, 4]; however, a later meta-analysis [5] revealed no consistent correlation across studies. Here, we found that variable genes were actually substantially *less* likely to be correlated to the B/F ratio (27%, *q ≤* 0.05) than either invariable (71%) or non-significantly-associated (55%) genes. These results parallel what we observe when we correlate gene family abundances with the α-diversity of observed bacterial species. We estimated α-diversity using the Shannon index, which is low when the distribution of species abundance is highly skewed, and high when there are many species of even abundance. Only 17% of significantly variable genes correlate significantly to the Shannon diversity (*q ≤* 0.05), versus 45% of significantly invariable and 26% of non-significant genes. We therefore conclude that bacterial and archaeal gene families identified as variable in this study are less likely to be associated with average genome size, B/F ratio, or α-diversity

When examining the PD-stratified gene families, we noticed that the variable/high-PD gene set was also enriched for gene families described as “hypothetical” in the KEGG Orthology database; hypothetical gene families were also observed in the invariable/low-PD set, but they were statistically depleted (see main text). We were interested in whether these conserved-yet-variable hypothetical gene families could be acting as markers for minor phyla. Indeed, out of 81 genes in this group, 44 were significantly associated with Proteobacterial abundance (*q ≤* 0.05 by the above Kendall’s partial τ test) and 13 were associated with Actinobacteria at the same threshold. However, 5 and 7 each were associated with Firmicutes and Bacteroidetes, indicating that even the major phyla of the human gut vary with respect to certain as-yet-uncharacterized functions.

